# Fate patterns originate through structural relations between cell and supracellular levels of organization

**DOI:** 10.1101/2025.10.24.683925

**Authors:** Clint S. Ko, Ruonan Chen, Nina T. Magid, Katharine Courtemanche, Pearson W. Miller, Alan R. Rodrigues, Amy E. Shyer

## Abstract

How patterns of cell state emerge across a tissue field is a fundamental question brought into renewed focus by spatial omics tools that map molecular states onto tissue organization. Here, we investigate how a field of limb progenitor mesenchyme transforms into distinct, adjacent cartilage and soft tissue compartments. We find that mesenchymal tissue fields self-organize their own differentiation through co-constitutive relationships between cell and supracellular structures, which produce cell-ECM or cell-cell-based supracellular cues that canalize cartilage or soft tissue cell fate change, respectively. At the tissue level, bifurcation in intrinsically generated supracellular structures guides the specification of tissue compartment size. We find that Wnt secreted from neighboring epithelial tissue influences mesenchymal cell fate and patterning by functioning as a modulator of cell-supracellular structural relations. Taken together, our results provide insight into how mesenchymal self-organization interfaces with epithelial signaling to enable a tissue compartmentalization process that initiates the skeleton.

## Introduction

Recent spatial omics techniques, which map wide-scale molecular activity to native tissue contexts, have underscored that cellular states are tightly aligned with structural features of tissue architecture^1-3^. Such characterizations, which link patterns across length scales, highlight the need to consider the causal relationships across levels of organization. In contexts spanning development, disease, and aging^2^, causal understanding is predominantly guided by models from developmental biology^4,5^, where differences in signaling molecule concentration across a tissue provide positional information mediated through subcellular mechanisms, which then templates the formation of tissue structures^6-13^.

Recently, the reverse causal direction, wherein tissue-scale mechanical and material properties feed back down to alter cell state, has been increasingly considered^14-16^. These causal relationships between cell states and tissue mechanics form the basis of recent mechanochemical frameworks^17-27^. Despite these insights, the full potential of cross-scale regulation in living systems is still in its infancy. Along these lines, in living systems, a particular class of scale-crossing coupling has been proposed where a macro-scale process and a micro-scale process are each reciprocally required for and generated by the other^28-30^. When these processes occur within a given entity (e.g. a tissue field and component cells), such a dynamic has the potential to imbue the system with the capacity to autonomously drive its own transformation (e.g. tissue differentiation)^30-33^. Such a co-constitution where the parts are themselves transformed by their participation in the whole they build has been proposed as a defining feature of self-organization in biological hierarchies as compared to inorganic systems^34-36^. For example, water molecules do not fundamentally change identity within a snowflake as it forms, nor do Lego blocks transform as they are incorporated into a structure.

Here, we focus on limb progenitor tissue fields to investigate whether co-constitutive relationships between the cell and supracellular level^37^ may clarify how coordinated patterns of cell state and tissue structure emerge. The early limb bud is a classic embryological system that has been subjected to extensive molecular, genetic, and cell biological analysis^38-45^. However, despite decades of study through patterning frameworks based on graded molecular control^46^, how mesenchymal progenitor fields initiate neighboring skeletal and soft tissue compartments has yet to be fully elucidated^47-50^, suggesting that a consideration of the relations between cell and supracellular processes could be fruitful. To investigate the possibility that cell-supracellular based self-organization leads to the initiation of spatial patterns of cellular differentiation, we optimized an *ex vivo* platform, called organite culture, using primary progenitor cells from the epithelial and mesenchymal tissues of the embryonic chicken limb. The organite platform reconstitutes a minimal version of the limb tissue ecosystem and captures the process whereby a field of homogenous mesenchymal progenitor cells, in the presence of secretory epithelial tissue, compartmentalizes into regions fated to become cartilage and soft tissue.

Combining the organite assay with cross-scale static and live-cell imaging allowed us to uncover that mesenchymal field self-organization arises through a strong coupling between cell-level and supracellular-level structures. Using experiment and mathematical modeling, we present a mechanism whereby cell-supracellular structural relationships serve as the central organizing system that mediates fate specification, processing of epithelial signals, and generation of precise compartment boundaries as well as pattern length scale. We find that both cartilage and soft tissue fate are initiated by distinct supracellular structures that are generated from within each collective, rather than from an external boundary, microenvironment, or substrate. In addition, recognizing the co-constitutive nature of cell- and supracellular-level structures allowed us to identify a functional role for signaling molecules secreted by the epithelium that differs from those based on graded molecular control.

### Reconstitution of tissue compartment patterning in an *ex vivo* primary progenitor assay

In the nascent limb bud (HH 21-23), the limb mesenchymal progenitor field compartmentalizes into adjacent core and peripheral domains marked by Sox9 and Twist expression, respectively, to initiate the skeleton (Figure 1A)^51-52^. Twist-positive cells, indicative of cells fated to become soft tissue, form adjacent to the ectoderm, surrounded by a field of Sox9-positive cells, indicative of cells fated to become cartilage. We observed that these patterns of gene expression state at the cell level correspond with distinct structures at the supracellular level. The Sox9-positive core region is associated with a cell-ECM network structure^53-54^, whereas the Twist-positive peripheral region is associated with high cell density and N-cadherin (Ncad) expression^55^, indicative of a cell-cell network structure (Figures 1B-1C, and S1A). Thus, limb mesenchyme compartmentalization is associated with the formation of novel and coincident properties at both cell and supracellular levels. We sought to understand how these multi-level patterns emerge and whether cross-level interactions could be the cause of such coincident patterns necessary for templating the emerging limb skeleton.

**Figure 1.**
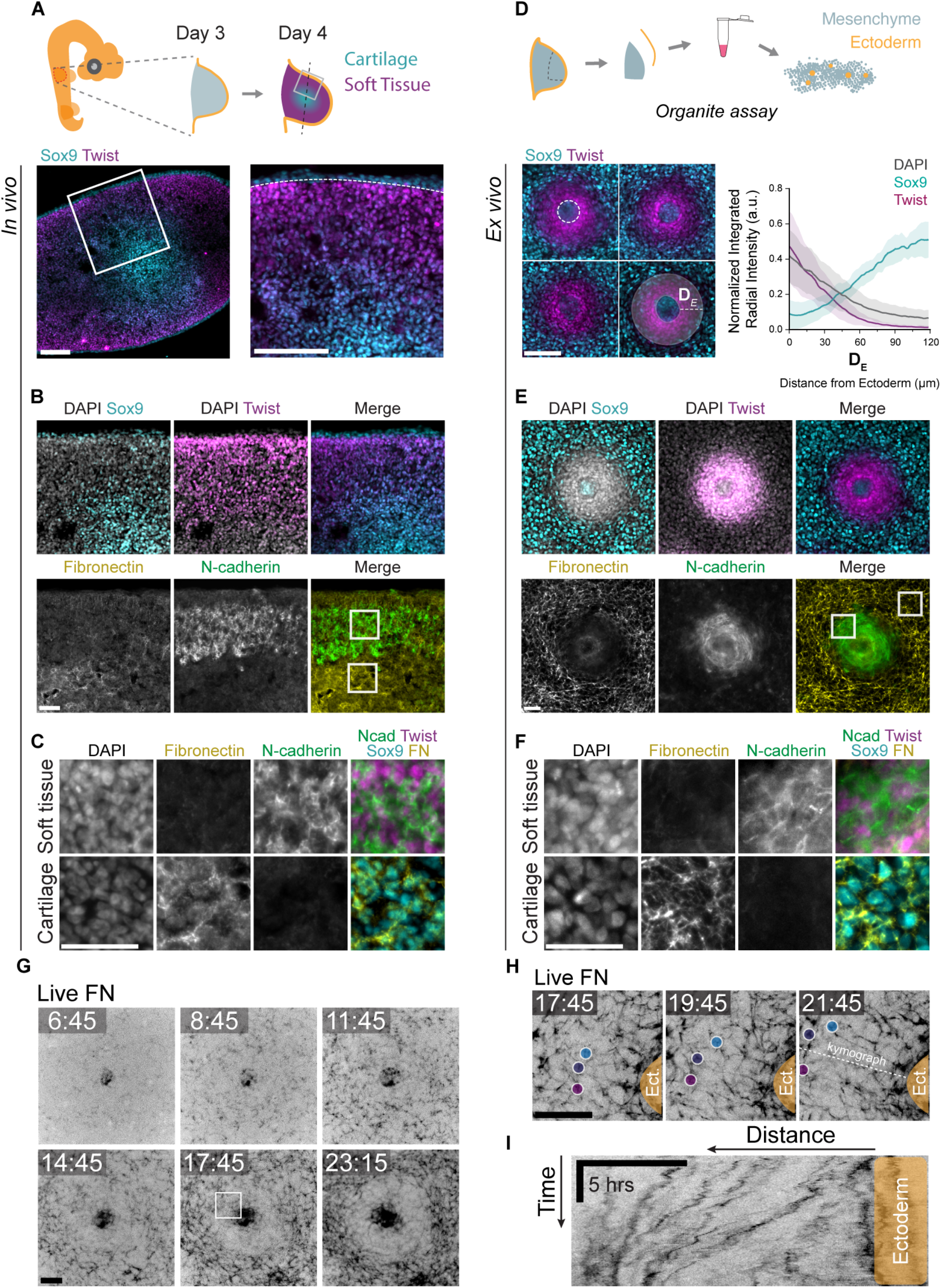
The organite platform reconstitutes cell state and supracellular structure patterns of cartilage and soft tissue fated fields. **(A)** Top, schematic showing orientation of transverse sections through limb buds. Bottom, transverse sections of E4.5 limb buds stained for Sox9 and Twist. White dashed lines are drawn at the boundary between the ectoderm (which is also Sox9+) and mesenchyme. **(B)** Sections of E4.5 limb buds stained for DAPI, Sox9, and Twist (top) or FN and Ncad (bottom). Scale bar, 40 µm. **(C)** Crops of regions in white boxes in 1B. Scale bar, 40 µm. **(D)** Top, schematic of *ex vivo* progenitor cell platform (organite). Bottom left, replicates of organites stained for Sox9 and Twist. Bottom right, Radial intensity profiles of normalized mean integrated intensities of DAPI (grey), Sox9 (cyan), and Twist (magenta) measured from the edge of the ectodermal tissue, D_E_ (see overlay in left images). Error bars represent mean ± SD. White dashed lines are drawn around the ectoderm. **(E)** Organites stained for DAPI, Sox9, and Twist (top) or FN and Ncad (bottom). Scale bar, 40 µm. **(F)** Crops of regions in white boxes in 1E. Scale bar, 40 µm. **(G)** Montage from a live imaging experiment visualizing fibronectin in organites (Movie S3). Scale bar, 50 μm. **(H)** Montage of region similar to that highlighted by white box in 1G where FN foci are marked with colored circles and tracked over time, showing displacement away from the ectoderm (Movie S4). Scale bar, 50 μm. **(I)** Kymograph from white dashed line in 1H. Horizontal scale bar (distance), 50 µm. Vertical scale bar (time), 5 hours. Scale bars are 100 µm unless otherwise noted.

Doing so requires access to spatiotemporal dynamics across cellular and supracellular levels of organization, which remains technically challenging to achieve *in vivo*. To obtain such access, we developed an *ex vivo*, primary progenitor cell-based platform that successfully recapitulates the tissue compartmentalization pattern observed in the limb in an inverted geometry. Specifically, to capture both the inductive capacity of epithelial tissue^56-57^ and the intrinsic self-organizing capacity of mesenchyme^19,58-59^, a dense disc of stacked cells composed of a defined ratio of dissociated progenitor epithelial (ectodermal) and mesenchymal cells from the embryonic day four avian limb bud is seeded and allowed to spontaneously arrange into islands of ectodermal tissue surrounded by a field of mesenchyme (Figures 1D, S1B-S1C, and Movie S1). We call this platform the organite assay for two reasons. First, the suffix - ite, denoting the constituents of a place, marks our use of primary progenitors, in contrast to many *ex vivo* organ models^60-62^ that utilize adult stem cells, which serve regenerative functions in established organ structures. The progenitor cells used in this platform are the cells that initially establish tissue organization within organs and, therefore, are ideally suited for the study of organ creation. Second, the - ite also stands for <u>i</u>nteracting <u>t</u>issues *<u>e</u>x vivo*. The inclusion of progenitors from both epithelial and mesenchymal types dispenses with the need to supplement with exogenous matrix in place of the mesenchyme or add exogenous differentiation cues in lieu of the epithelium. Further, because both tissues are dissociated and, therefore, any molecular prepattern or template is reset, the system captures the pattern from its inception.

After 24 hours of culture, the organite platform reproduced a stereotyped tissue differentiation pattern in the mesenchymal population in which a field of Twist-positive cells formed adjacent to the ectoderm, surrounded by a field of Sox9-positive cells (Figures 1D-1E). In addition to recapitulating cell fate patterns, the organite platform recapitulates supracellular structural patterns. Specifically, the Twist-positive region adjacent to the epithelium recapitulates the high cell density and Ncad expression that we observed in the peripheral limb bud domain, with densely packed, elongated cells (Figures 1E-1F, S1D, and Movie S2). In the Sox9-positive region, a well-developed cell-ECM network is observed, marked by fibronectin (FN) (Figures 1E-1F) as well as tenascin^63-64^ and thrombospondin^65-66^ (Figure S1E). We also observe the established cartilage fate marker, type II collagen^67-69^, specific to the Sox9-expressing region in the organite (Figure S1E).

Having validated that the organite recapitulates key cell state and supracellular features of the pattern, we utilized the organite platform to spatiotemporally observe the pattern-forming process. We observe that in the early phase (0-9 hours) of organite culture, fibronectin is produced by the ectoderm^70^ (Figure S1F) and expressed across the mesenchyme (Figure 1G and Movie S3). Strikingly, after approximately 10 hours, we observe an outward movement of fibronectin away from the ectoderm (Figures 1G-1I, Movie S3, and Movie S4), producing a low-ECM zone in the putative soft tissue domain and a fibronectin-cell network specific to the cartilage-fated domain. We observe that fate marker expression evolves concurrently with, rather than prior to, fibronectin rearrangement (Figure S1G), leaving open the possibility that supracellular cell-ECM structure, rather than being a consequence of fate patterning, could be the cause of Sox9 expression. We therefore investigated the link between supracellular cell-ECM structure and Sox9 expression.

### Supracellular cell-ECM structure is sensed through actin structure to initiate cartilage fate

When primary limb mesenchyme progenitors are cultured in the absence of ectoderm or other extrinsic tissue layers or signals, the majority of mesenchyme adopts cartilage fate^71^ (Figures 2A and 2C), making this a valuable system to test the hypothesis that cell-ECM supracellular structure could be the cue for Sox9 expression. Consistent with a role for supracellular structure in Sox9 expression, when cell density is modulated in mesenchyme-only (MO) culture, we found that a particular density threshold is required for Sox9 expression (Figures 2B and 2D). To investigate whether the link between density and Sox9 involves the formation of a cell-ECM network, we performed cross-scale spatiotemporal analysis in MO culture. We found that supracellular scale changes in cell-ECM network structure, as measured by FN and coherence of focal adhesion components enriched at cell interfaces, were spatiotemporally concomitant with Sox9 marker expression (Figures 2E, S2A-S2B, and Movie S5).

**Figure 2.**
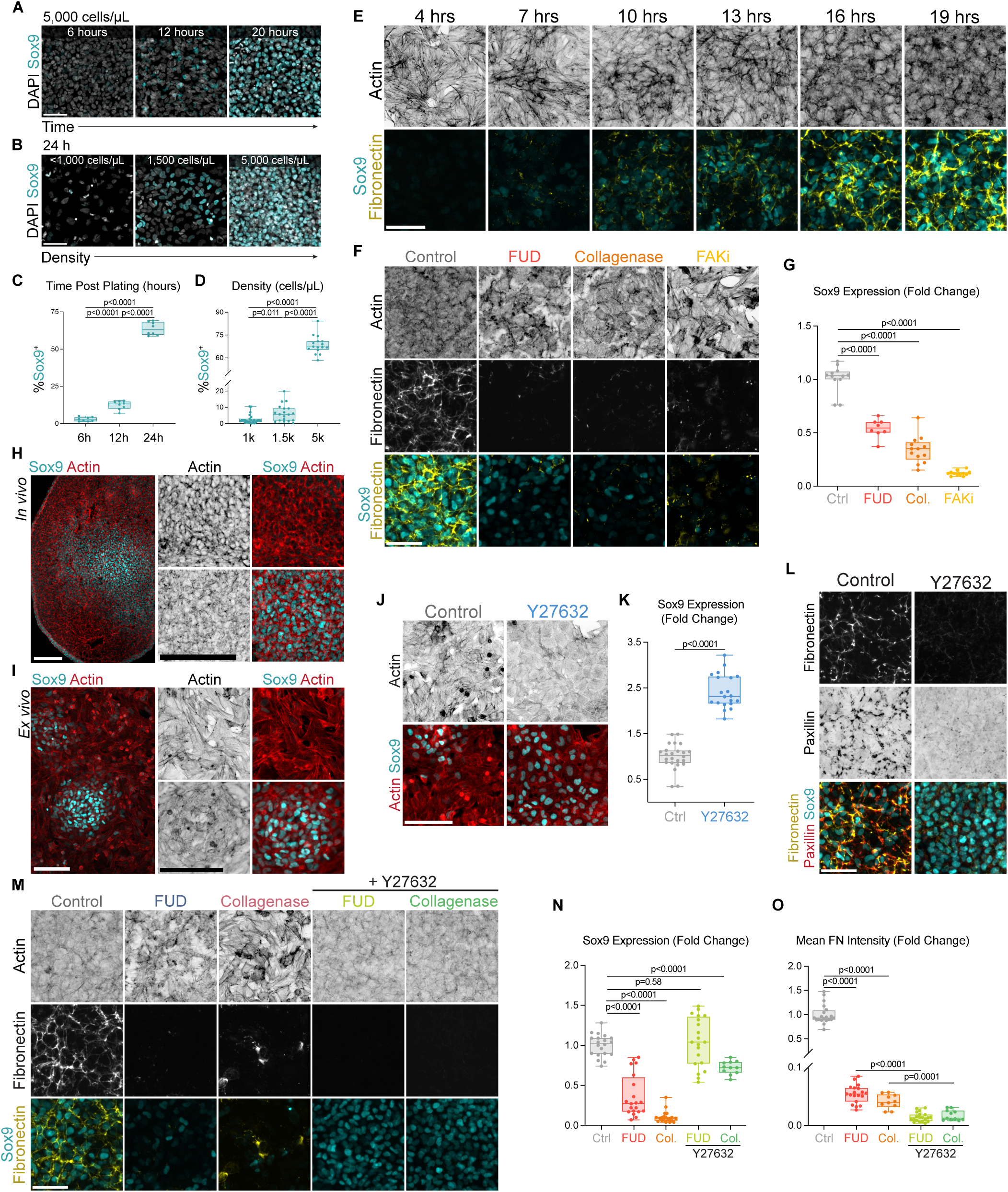
Supracellular structure initiates cartilage fate through changes in actin architecture. **(A)** Sox9 levels increase over time until most limb progenitor cells adopt cartilage fate by 24 hours of culture. Scale bar, 50 µm **(B)** DAPI and Sox9 in 24-hour *ex vivo* cultures plated at different densities show Sox9 activation is density-dependent. **(C)** Quantification of %Sox9+ area at different culture times. **(D)** Quantification of %Sox9+ area in cultures plated at different densities. **(E)** Time course of mesenchyme-only (MO) cultures stained for Actin, Sox9, and FN revealing changes in supracellular structure associated with Sox9 expression. **(F)** Actin, FN, and Sox9 in 24-hour MO cultures treated with FUD (0.5 µM), collagenase (0.5 µg/mL), and FAKi (20 µM). **(G)** Quantification of Sox9 intensity levels (fold change) after treatment with FUD, collagenase, and FAKi. **(H)** E5 limb bud section stained for DAPI, Sox9, and Actin shows a pattern of more diffuse actin fibers in Sox9-positive cells as compared to Sox9-negative cells. Scale bars, 100 µm. **(I)** MO culture stained for actin and Sox9. Scale bars, 100 μm. **(J)** Sox9 and actin in MO cultures treated with Y27632 (10 µM). Scale bar, 100 µm. **(K)** Quantification of % Sox9+ area. **(L)** MO cultures stained for FN, paxillin, and Sox9 demonstrate that Y27632 treatment leads to decreased cell-ECM engagement. **(M)** Actin, FN, and Sox9 in 24-hour MO cultures treated with FUD, collagenase, and Y27632. **(N)** Quantification of Sox9 intensity (fold change). **(O)** Quantification of mean FN intensity (fold change). Scale bars are 50 µm unless otherwise noted.

We investigated whether such supracellular structure is necessary for Sox9 expression by disrupting cell-ECM network formation through three distinct treatments. Indeed, inhibition of FN polymerization through the FUD peptide^72^, degradation of ECM through mild collagenase treatment, or inhibition of focal adhesion kinase (FAK) (FAK inhibitor 14, FAKi) to affect focal adhesion stability and dynamics^73-74^ each lead to a significant reduction or loss of Sox9 expression (Figures 2F-2G), accompanied by gross alterations in levels or structure of the cell-ECM network as assessed by ECM molecule and focal adhesion component localization (Figures S2C-S2E). Thus, Sox9 gene expression depends on information generated through the structure of emergent cell-ECM networks.

Next, we sought to understand how the emergence of supracellular structure could influence cell-level decision-making to activate Sox9 expression. In our cross-scale structural analysis of ECM at the supracellular scale (Figure 2E), we observed that the development of an ECM network corresponds with evolving actin architecture at the cell level. Further, we observed an association between Sox9 expression and cytoskeletal architecture *in vivo* (Figure 2H) and in MO culture (Figure 2I), where Sox9 is expressed in cells with diffuse cortical actin and reduced cable/fiber formation or bundling. Loss of an ECM network drives an increase in actin cable formation and bundling (Figure 2F), indicating that cytoskeletal architecture is governed by supracellular cell-ECM network structure.

To test whether cytoskeletal architecture mediates the effect of supracellular structural changes on Sox9 expression, we induced a diffuse cortical actin architecture using the Rho-associated coiled-coil kinase (ROCK) inhibitor Y-27632^75-76^. At plating densities that normally result in low Sox9 expression, Y-27632 treatment leads to near-ubiquitous Sox9 expression (Figures 2J-2K), consistent with observations in single-cell studies^77-79^. Notably, this rescue of Sox9 expression occurs in the absence of matrix and focal adhesion network formation (Figure 2L). In further confirmation that such activation can occur in the absence of cell-ECM structure, we inhibited the formation of a cell-ECM structure in MO culture through FUD or mild collagenase treatment and found that we rescued Sox9 expression by inhibiting actin cable formation with Y-27632 (Figures 2M-2O, and S2F). These results indicate that, provided that the cytoskeletal architecture is in the proper configuration, Sox9 activation does not require cell-ECM supracellular structure.

Together, these results place cytoskeletal architecture downstream of supracellular cell-ECM structure, where it serves as an intermediary through which supracellular cues enable Sox9 activation. Further, these results support a model where focal adhesions and fibronectin are not functioning at the single-cell level to enable Sox9, but rather that cell-ECM structures at the supracellular scale impose cell fate across the entire collection of cells by inducing a particular cytoskeletal architecture.

### Epithelial Wnt activates soft tissue fate by stimulating emergent cell-cell supracellular structure

Established ectodermal signals are known to prevent cartilage formation near the ectoderm and enable soft tissue fate^40,51,80-83^. Given that a cell-ECM supracellular structure guides cartilage fate initiation in the mesenchyme, we investigated whether secreted ectodermal cues, such as Wnt (Wnt3a)^84^, bias progenitors away from cartilage and toward soft tissue by acting on that same supracellular structure. We confirmed that, in MO culture, Wnt treatment leads to a loss of Sox9 expression (Figure 3A). Notably, we also observed the disruption of cell-ECM network formation, as assessed by paxillin (Figure 3A) and FN (Figure 3B and Movie S6). In line with the model that Wnt limits cell-ECM engagement rather than simply blocking ECM production, we found that exogenous recombinant FN added to Wnt-treated MO cultures did not restore cell-ECM network formation or Sox9 to the levels of FN-supplemented control cultures (Figures 3C-3F).

**Figure 3.**
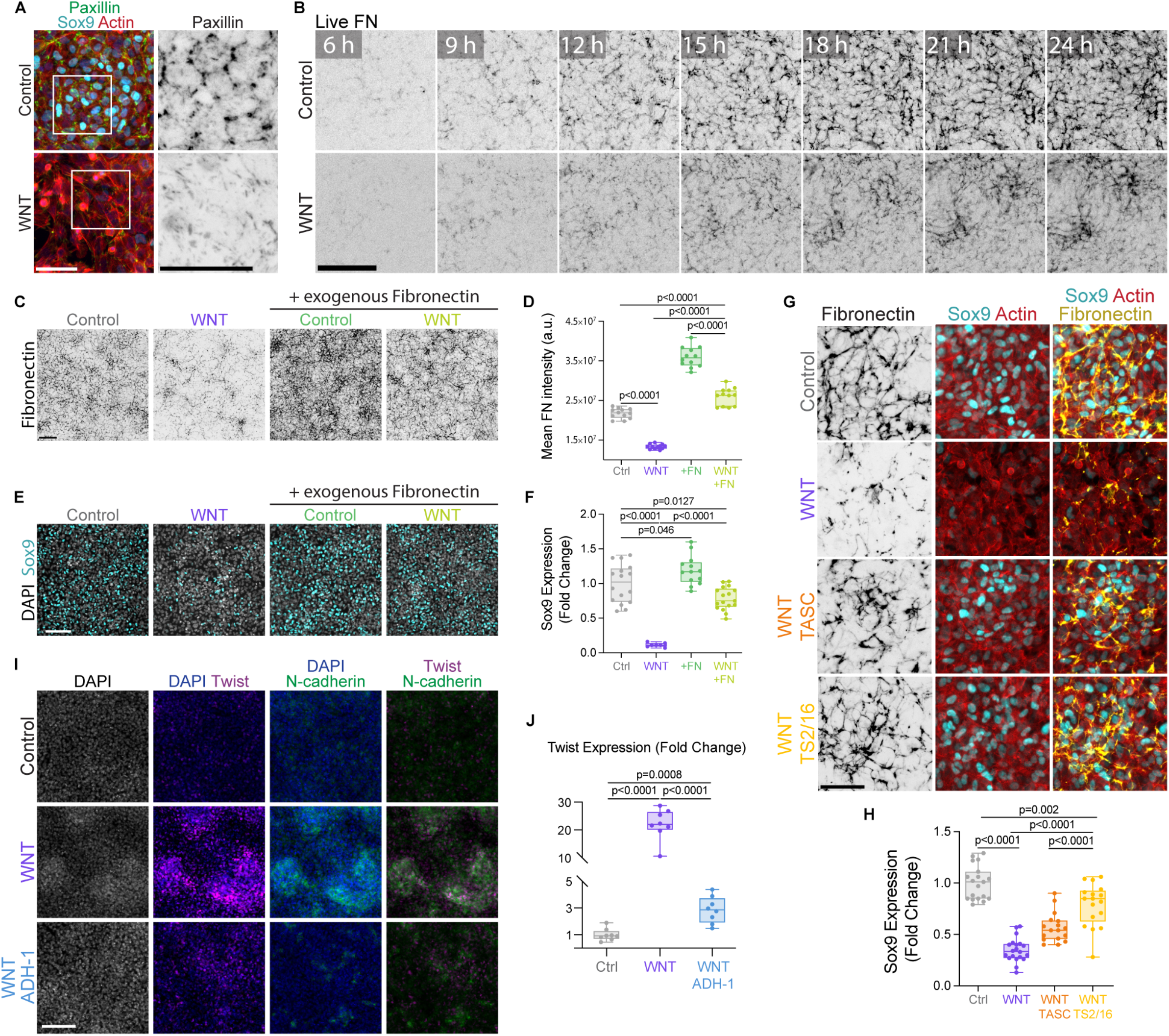
Wnt activates soft tissue fate by suppressing cell-ECM engagement and promoting cell-cell adhesion. **(A)** Wnt treatment disrupts cell-ECM structure formation, reducing paxillin levels, to inhibit Sox9. Control and WNT-treated (150 ng/µL) MO cultures stained for paxillin, Sox9, and actin. Right images show crops of regions highlighted by white boxes on the left set of images. Scale bar, 50 µm. **(B)** Montage from live imaging of FN in control and Wnt-treated (150 ng/mL) MO cultures (Movie S6). Scale bar, 50 μm. **(C)** FN staining of control and Wnt-treated MO cultures supplemented with exogenous recombinant FN (5 μg/mL). **(D)** Mean FN intensity comparison of 3C. **(E)** Sox9 staining of control and Wnt-treated MO cultures supplemented with exogenous recombinant FN. **(F)** Quantification of Sox9 intensity (fold change) of 3E. **(G)** MO cultures treated with WNT and β1 integrin activating antibodies, TASC (20 µg/mL) and TS2/16 (20 µg/mL), stained for FN, Sox9, and actin. Integrin activation rescues cell-ECM structure and Sox9. Scale bar, 50 µm. **(H)** Quantification of changes in Sox9 levels for 3G. **(I)** MO cultures treated with WNT alone or in addition to the Ncad inhibitor ADH-1(.25 mg/mL) stained for DAPI, Ncad, and Twist show WNT induces aggregation and Twist activation through promoting a cell-cell adhesion-based structure. **(J)** Quantification of fold change in Twist intensity normalized to DAPI. Scale bars are 100 µm unless otherwise noted.

To test whether Wnt inhibits Sox9 through reducing the cell-ECM supracellular structure that we found is required for Sox9 expression, we amplified cell-ECM engagement through integrin β1-activating antibodies TASC^85^ and TS2/16^86^ in cells treated with Wnt and assayed for Sox9 expression. Under these conditions, cell-ECM supracellular structure was restored, and Wnt could not properly repress Sox9 (Figures 3G-3H), indicating that the effect of Wnt on Sox9 expression requires a decrease in cell-ECM engagement.

Given these results, and motivated by the elevated cell density and Ncad expression that mark the soft tissue domain, we asked whether Wnt promotes soft tissue fate by driving an alternative supracellular structure that is based on cell-cell adhesion. In MO culture, we found that Wnt enables Twist expression, drives increased Ncad expression, and increases cell aggregation (Figures 3I-3J, S3A, and S3B), indicating that Wnt biases limb progenitor collectives to form a dense cell-cell adhesion-based supracellular structure rather than a cell-ECM-based structure. Furthermore, we found that Wnt cannot activate Twist or increase cell packing when Ncad function is impeded through competitive binding (ADH-1)^87-88^ (Figures 3I-3J), indicating that Wnt influences Twist activation through changes in cell-cell relations and the resulting supracellular structure, rather than in a direct, cell-autonomous manner.

### A scale-crossing co-constitutive causal logic governs cartilage-initiating tissue structure

Given our finding that limb progenitor cell fate initiation is the result of distinct supracellular structures, we sought to determine how these structures form. In the case of limb progenitors initiating cartilage fate through cell-ECM tissue structure, it is notable that this transition can occur without the addition of exogenous matrix or specific growth factors. Instead, the field of progenitors organizes this structure from within, thereby distinguishing this system from those in which cells are placed upon and respond to an exogenous matrix substrate^89-94^.

We investigated the features that enable the generation of structure from within the collective, and observed that intracellular calcium (Ca^2+^), a means by which cells collectively coordinate ECM assembly^59,95-98^, is necessary for the development of the cell-ECM network that enables Sox9 expression. Specifically, decreasing intracellular calcium levels with BAPTA-AM disrupts cell-ECM network formation and Sox9 activation in a dose-dependent manner (Figures 4A-4B, 4D). We further confirmed the necessity of calcium signaling by inhibiting the calcium sensor and effector calmodulin^99^ (Figure S4A). At the highest levels of inhibition with BAPTA-AM, when cell-ECM network formation is completely blocked, exogenous FN is unable to rescue network formation, suggesting that Ca^2+^ is not required for supplying or producing fibronectin but for its assembly and organization into the appropriate supracellular cell-ECM structure (Figures 4C-4D). Further, at high levels of inhibition, supplemental FN does not lead to increased Sox9 (Figures 4B-4C), suggesting that FN promotes Sox9 not as a molecular signaling factor, but as part of a supracellular network formed through calcium-dependent processes.

**Figure 4.**
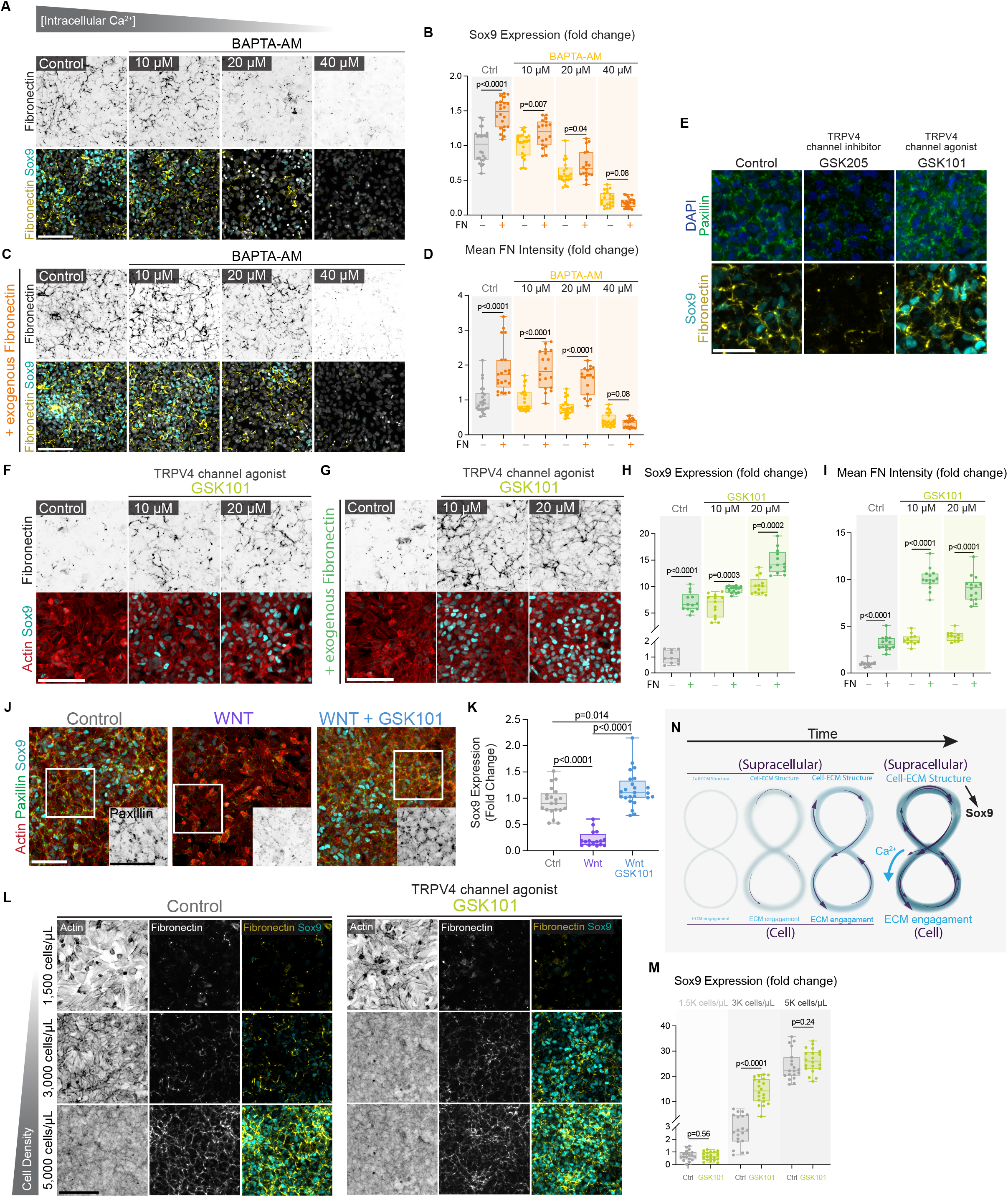
Cross-scale structural relations generate the cell-ECM structure that initiates cartilage fate. **(A)** Medium-high density MO cultures treated at a range of concentrations (10-40 µM) of the intracellular calcium chelator BAPTA-AM stained for FN and Sox9. **(B)** Quantification of changes in Sox9 levels **(C)** Control and BAPTA-AM-treated MO cultures with the addition of exogenous recombinant FN (5 μg/mL). Sox9 levels for cultures with exogenous FN are shown in orange in 4B. **(D)** Mean FN intensity in MO cultures treated with BAPTA-AM, with or without exogenous FN. **(E)** MO cultures treated with a TRPV4 antagonist, GSK205 (50 µM), or agonist, GSK101 (15 µM), stained for DAPI, paxillin, Sox9, and FN. Scale bar, 50 µm. **(F)** Medium-low density MO cultures stained for FN, actin, and Sox9 treated with increasing concentrations of the TRPV4 agonist (10 – 20µM), driving greater cell-ECM engagement. **(G)** Same culture conditions as 4F but with the addition of exogenous FN, showing enhanced incorporation of FN into a network and greater Sox9 activation. **(H)** Quantification of Sox9 levels (fold change) for 4F and 4G. **(I)** Quantification of FN intensity (fold change) for 4F and 4G. **(J)** Treatment with TRPV4 agonist GSK101 (15 µM) rescues cell-ECM engagement and cellular cohesion in WNT-treated (150 ng/mL) MO cultures to rescue Sox9. Insets show paxillin staining in the white boxes. Scale bars, 75 µm. **(K)** %Sox9+ area of 3A. **** p < .0001. **(L)** MO cultures plated at a range of cell densities (1,500 to 5,000 cells/µL) reveal that GSK101 treatment (20 µM) lowers the threshold for cell-ECM network formation but fails to rescue Sox9 expression at the lowest densities. MO cultures stained for actin, FN, and Sox9. **(M)** Quantification of changes in Sox9 levels in 4L. Scale bars are 100 µm unless otherwise noted. **(N)** Schematic depicting a cross-scale co-constitution that generates a cell-ECM supracellular structure to initiate cartilage fate. Supracellular cell-ECM structure triggers structure-sensitive TRPV4 channels to enable Ca^2+^-dependent cell-ECM engagement, just as cell-ECM engagement enables the supracellular cell-ECM structure.

Under moderate calcium inhibition (10 or 20 µM BAPTA-AM), Sox9 expression and FN network formation are reduced (Figures 4A-4D). If the inhibitor suppressed Sox9 cell-autonomously, adding exogenous FN should not rescue Sox9, since intracellular calcium is equally blocked with or without it. Instead, under moderate calcium inhibition, supplying cultures with additional FN increases both cell-ECM network formation and Sox9 expression (Figures 4B and 4D), further evidencing that intracellular calcium enables Sox9 activation through the formation of a supracellular cell-ECM structure, rather than through a cell-autonomous mechanism. Testing for gain of function, we observed that strongly enhancing calcium influx through treatment with Bay K8644^100^, an agonist for L-type calcium channels^101^, leads to a dose-dependent increase in FN network formation and Sox9 expression, both with and without supplemental FN (Figures S4B-S4E).

Given the critical role of intracellular calcium in the development of the cell-ECM supracellular structure, we sought to determine how this occurs at the cellular level. We investigated candidates and found that the mechano-responsive ion channel TRPV4 is essential for cell-ECM network formation. TRPV4 inhibition reduced focal adhesions and cell-ECM supracellular structure (Figure 4E). As predicted from its dependency on such structure, Sox9 expression was also reduced. Conversely, when TRPV4 channels were constitutively activated by an agonist, focal adhesions, cell-ECM network formation, and Sox9 were enhanced (Figure 4E), identifying TRPV4 channels as key mediators of the intracellular Ca^2+^ influx necessary for cell-ECM network formation and subsequent Sox9 activation. We supplied exogenous FN to cultures alongside increasing concentrations of TRPV4 agonist and observed that FN incorporation and Sox9 activation scaled with agonist concentration (Figures 4F-4I), providing additional evidence that TRPV4 activity drives cell-ECM engagement and supracellular cell-ECM structure formation.

We assayed whether TRPV4 activity can rescue experimentally attenuated cell-ECM engagement. Indeed, when MO cultures were treated with FAKi (Figures S4F-S4G) or Wnt (Figures 4J-4K), ectopic TRPV4 activation rescued the formation of a cell-ECM structure and Sox9 expression, further substantiating that TRPV4 activity is sufficient to induce cell-ECM engagement. To validate this through orthogonal means, we attenuated cell-ECM engagement by reducing plating density. At intermediate densities, below the threshold for robust supracellular network formation but where cells remain close enough to contact one another, agonist-mediated TRPV4 activation induced cell-ECM network formation, together with the predicted actin architecture and Sox9 expression (Figures 4L-4M). These results reinforce that TRPV4 activity promotes cell-ECM engagement.

Strikingly, this rescue had a lower density limit. Specifically, at sparse densities where cells are too far apart to establish a connected structure, TRPV4 activation no longer rescued cell-ECM structure or Sox9 (Figures 4L-4M). TRPV4 activity, therefore, does not activate Sox9 expression in a cell-autonomous manner and depends on a minimal threshold of supracellular structure to trigger cell-ECM engagement. At high density, the effect of the agonist is lost (Figures 4L-4M), suggesting that the endogenous, structure-derived stimulus for TRPV4 is saturated in dense cultures. Stimulating L-type channels with Bay K8644 at high density further increases FN network formation and Sox9 levels, confirming that our high-density results reflect TRPV4 stimulus saturation rather than Sox9 readout saturation (Figures S4H-S4I). This is consistent with the well-established role of TRPV4 as a mechanotransducer that converts cell-ECM structure-derived membrane tension into Ca^2+^ influx^96-97,102-103^. Together with our finding that Ca^2+^ influx drives cell-ECM engagement, these results support a closed causal organization in which supracellular structure induces cell-ECM engagement through TRPV4 just as cell-ECM engagement generates supracellular structure (Figure 4N). In this model, through this co-constitutive relationship across scales, the collective self-organizes a cell-ECM supracellular structure that governs the cytoskeleton of the constituent cells to guide cells toward cartilage fate.

### A scale-crossing co-constitutive organization is stimulated by Wnt to enable soft tissue

We investigated whether cell-cell adhesion and densely packed supracellular structure are linked by a similar scale-crossing co-constitution, and whether Wnt acts on this relationship to drive soft tissue formation. It is well established that cell-cell adhesion leads to a dense, packed structure^104-106^. This is consistent with our own observation that, in MO and organite culture, Ncad expression is highest in packed regions, and that inhibiting Ncad function in MO culture reduces cell aggregation (Figures 1E-1F, 3I-3J). To test the less well-established causal direction, namely, whether packed supracellular structure prompts cell-cell adhesion, we varied density by modulating the initial plating cell number in MO cultures. In control cultures, although Ncad expression in the absence of Wnt is generally low, Ncad levels at the highest density increase disproportionately to cell number alone, suggesting that density itself can induce cell-cell adhesion (Figures 5A-5C). We observed this density-dependent effect even more strongly in the context of Wnt treatment. As density increased, Wnt-treated cultures showed a dramatic, nonlinear rise in Ncad levels that far exceeded what would be expected from increased cell number alone (Figure 5C). These results are consistent with a causal role for packed supracellular structure in promoting cell-cell adhesion. Along these lines, we found that while inhibition of Ncad through competitive binding (ADH-1) blocks the ability of Wnt to induce Ncad and Twist levels, we could rescue both Ncad and Twist in this context by increasing density (Figures 5D-5E). These results indicate that high cell density stimulates cell-cell adhesion and are consistent with a cross-scale co-constitution model where dense cell packing and cell-cell adhesion reciprocally generate and require one another.

**Figure 5.**
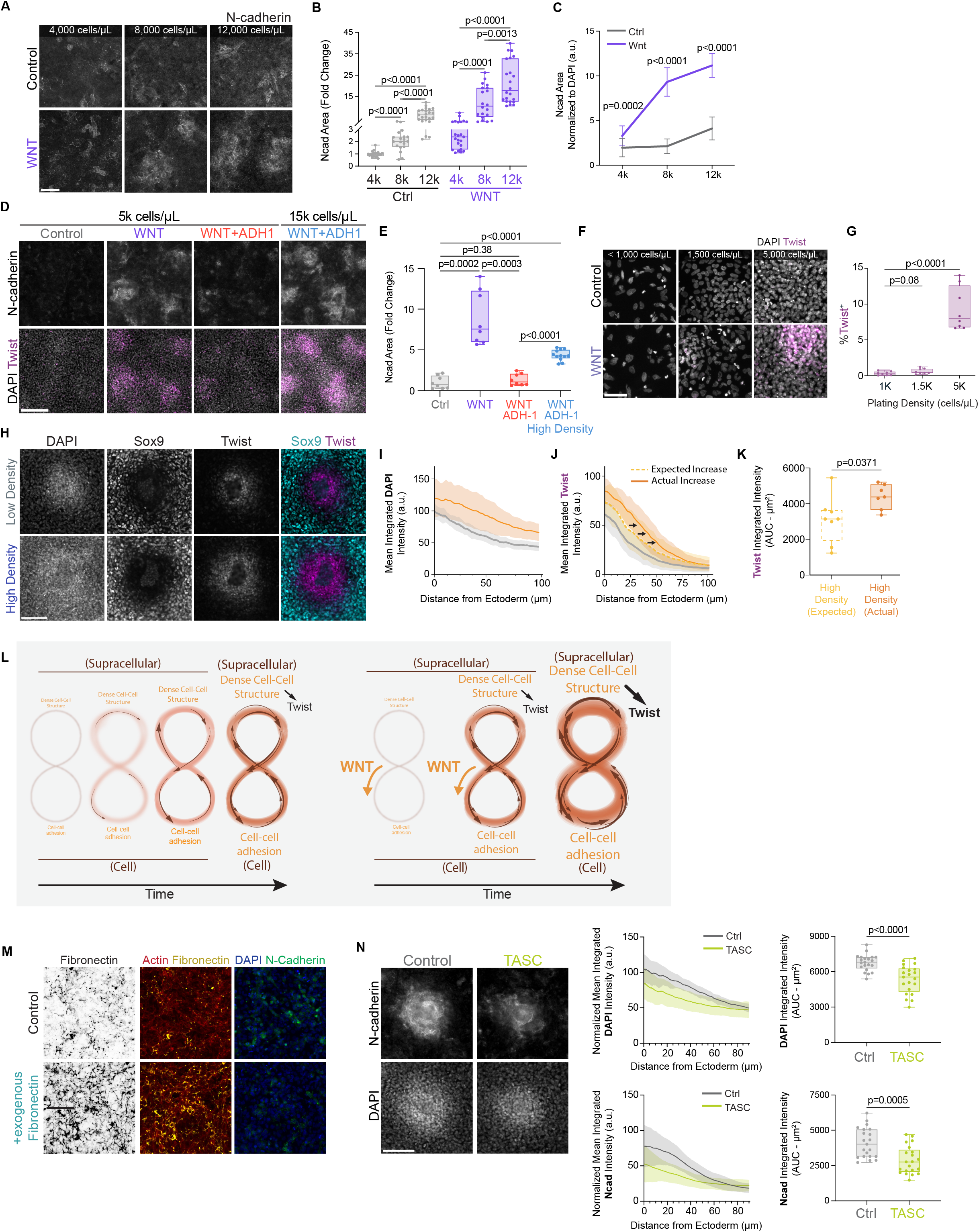
Co-constitution between cell-cell adhesion and dense supracellular packing is stimulated by Wnt to enable soft tissue fate. **(A)** Ncad in control or WNT-treated (150 ng/mL) MO cultures of different plating densities reveal WNT induces Ncad activation at the collective cell scale. Scale bar, 50 µm. **(B)** Quantification of Ncad area (fold change) **(C)** Ncad area quantification normalized to DAPI indicate a supralinear relationship where higher cell densities lead to greatly enhanced Ncad activation in WNT-treated cultures (150 ng/mL). **(D)** Plating at high cell densities rescues Twist and Ncad in MO cultures treated with WNT and the Ncad inhibitor ADH-1 (.25 mg/mL). Scale bar, 150 µm. **(E)** Quantification of changes in Ncad area. **(F)** Wnt treatment (150 ng/mL) in MO cultures at different plating densities shows induction of Twist downstream of WNT is density dependent. Scale bar, 50 µm. **(G)** Quantification of %Twist+ area of 3F. **(H)** Organites plated at low (5,000 cells/μL) or high cell density (20-40,000 cells/μL) stained for DAPI, Sox9, and Twist. **(I)** Radial intensity profile of normalized mean integrated intensities of DAPI in low- and high-density organites. **(J)** Radial intensity profile of normalized mean integrated intensities of Twist in low- and high-density organites. Dashed line represents the expected Twist signal based on DAPI intensity. **(K)** Area under the curve (AUC) comparison of expected and actual total integrated Twist intensities in high-density cultures. **(L)** Schematic depicting a cross-scale co-constitution that generates a packed cell-cell-based supracellular structure to activate Twist expression towards soft tissue fate. Dense cell packing enables cell-cell adhesion via Ncad, just as Ncad generates cell packing. Epithelial Wnt signal lowers the density threshold needed to enable increases in cell-cell adhesion. **(M)** MO cultures treated with exogenous recombinant FN display a decrease in Ncad. **(N)** (Left) DAPI and Ncad in control and TASC-treated (20 μg/mL) organites. (Right) Radial intensity profiles and AUC analysis of normalized mean integrated intensities of DAPI and Ncad. Scale bars are 100 µm unless otherwise noted. All error bars represent mean ± SD.

We next asked how such cross-scale reciprocity reframes the role of Wnt in enabling cell-cell adhesion. When starting cell density was lowered below a particular threshold, we observed only a slight increase in Ncad expression in Wnt-treated compared to control cultures, in contrast to the pronounced increase in Ncad observed at high density (Figures 5A-5C). These results further support that Wnt does not meaningfully increase Ncad levels in a cell-autonomous manner, but rather in a manner contingent on the density of the collective. Further, per-cell Ncad expression in low density Wnt-treated cultures is comparable to levels in control cultures at the highest plating density (Figure 5C), indicating that cells treated with Wnt adopt a cell-cell relational mode at lower densities than would otherwise be required. These results support a model where Wnt amplifies the co-constituting relationship by lowering the density threshold needed to cue cell-cell adhesion, which enables further increases in local cell packing. The result of this cross-scale co-constitution is a cell-cell adhesion-based supracellular structure that our results indicate promotes Twist rather than Sox9 expression.

We set out to probe whether such a cell-supracellular co-constitution mediates the effects of the Wnt signal on Twist expression. Having shown that cell-cell adhesion is necessary for Wnt to induce Twist (Figures 3I-3J), we tested the necessity of supracellular structure for Wnt to induce Twist. Indeed, when we varied density in MO cultures and treated with Wnt, we found that Wnt can only activate Twist in MO cultures when cells are above a particular density (Figures 5F-5G). We corroborated these results in organite culture, where the ectoderm supplies Wnt. To do so, we created a highly packed mesenchyme through increasing the starting mesenchymal cell density and observed increased Twist expression (Figure 5H). This increase in expression is amplified in a nonlinear manner as compared to the increase in cell number (Figures 5I-5K), indicating that the capacity for Wnt to enable cell-cell adhesion and subsequent activation of Twist is contingent on supracellular structure, specifically densely packed cells (Figure 5L).

Given our results indicating that Wnt treatment increases cell-cell adhesion and reduces cell-ECM engagement, we investigated whether these relational modes are mutually suppressive. When MO cultures were supplemented with FN (Figure 5M) or integrins were ectopically activated (TASC) in MO and organite culture (Figures 5N and S5A), we observed a decrease in Ncad and cell packing. Thus, the addition of FN or activation of integrins, which each increase cell-ECM engagement, diminishes the cell-cell relational mode. We also examined whether increased cell-cell adhesion decreases cell-ECM engagement. In Wnt-treated MO cultures, even those with excess FN, incorporation of FN into a network declined with increasing cell density, where Ncad expression is highest (Figures S5B-S5C). As FN was held constant and in excess across all density conditions, this decline cannot be attributed to reduced FN availability, indicating that increased cell-cell packing and the ensuing increase in cell-cell adhesion suppress the cell-ECM mode. Together, these data are consistent with a mutually suppressive relationship between the two modes.

### (6) Competing cross-scale co-constitutive processes are sufficient to pattern tissue compartments without graded molecular control

Having established that supracellular structure enables cell fate, and that this structure emerges through cell-supracellular structural relations, we next asked whether such structural relations are also responsible for initiating skeletal patterning at the organ scale. Specifically, we asked whether two co-constitutive processes, each self-amplifying but mutually suppressive, can explain the spatial bifurcation of tissue fates. To do so, we developed a continuum model in which each cell’s intrinsic propensity to commit to a relational state (cell-cell versus cell-ECM) determines and is conditioned by the supracellular structure in which the cell resides as the system evolves. Local levels of engaged ECM (tension) set the threshold for the cell-ECM relational mode, while cell density sets the threshold for the cell-cell mode. While cell-ECM and cell-cell propensities are initially highly malleable, the temporal dynamics of cell commitment are treated as a bistable nonlinear switch (Figure 6A and Supplemental Note). Crucially, simulations do not begin with two distinct cell states that sort, as in phase separation based on differential adhesion^105-110^. Instead, the field starts nearly homogeneous, with only slight variation in cell-ECM and cell-cell propensity at length scales far below that of the final pattern.

**Figure 6.**
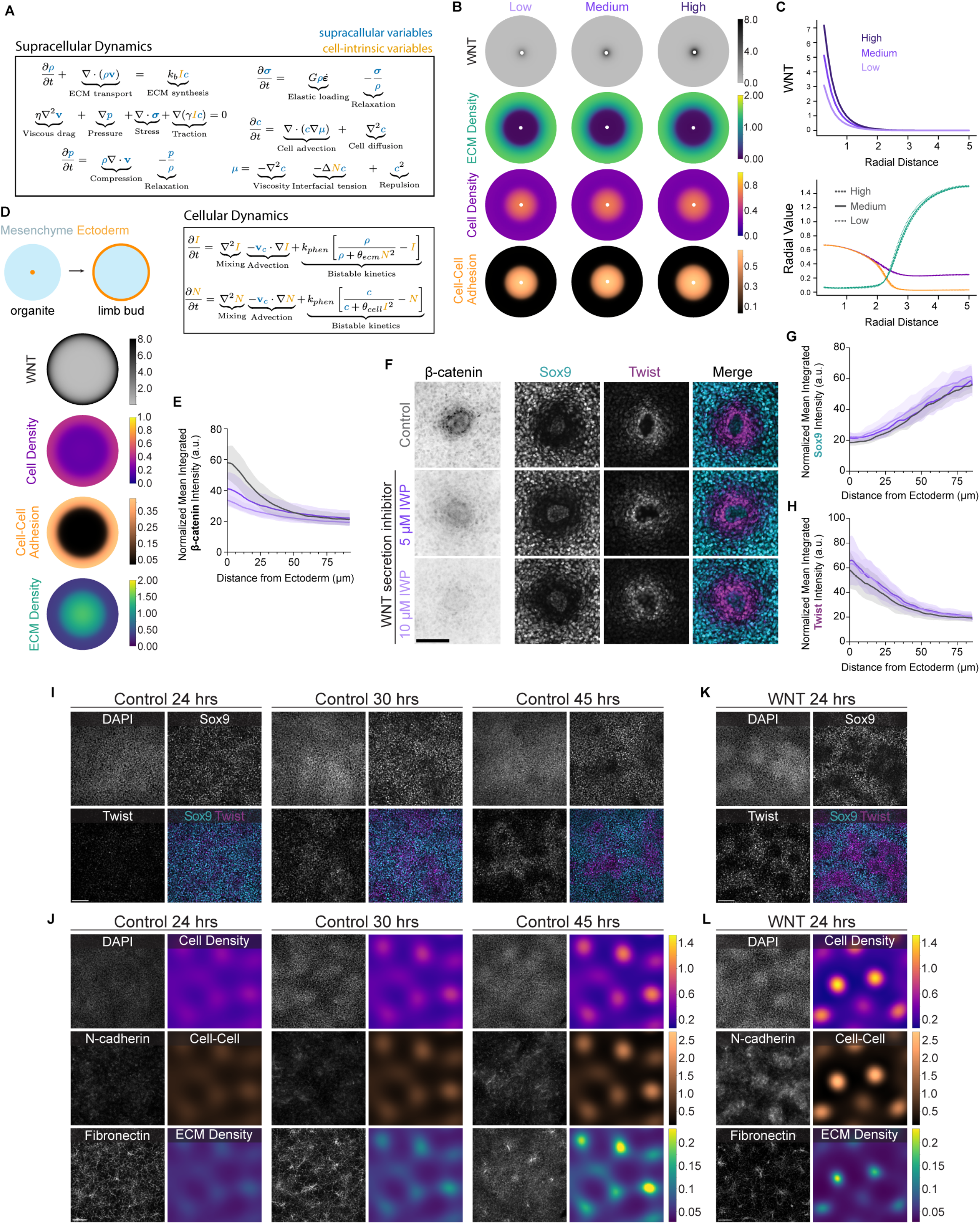
Cross-scale co-constitutive processes are sufficient to pattern tissue compartments without graded molecular control. **(A)** Key equations of mathematical model used throughout. See Supplemental Material for full details. **(B)** Simulations of organites on a disk domain of radius = 5 where Wnt is introduced from the boundary with ectodermal WNT factor = 3 (low), 5 (control), and 7 (high) reproduce *in vivo* tissue organization. Simulations show negligible differences in distributions of ECM, cells, and cell-cell adhesions, indicating that patterning length scales are set by bulk dynamics rather than through ectodermal signal strength. **(C)** Plots of radial cross-sections of key scalar fields from simulations **(D)** Simulation of cell and ECM densities on a disk domain of radius R = 5 where WNT is introduced from the boundary (see diagram on left) reproduces *in vivo* tissue organization. **(E)** Radial intensity profiles of normalized mean integrated intensities of β-catenin in organites indicate that treatment with an inhibitor of WNT secretion, IWP-2 (5 – 10 µM), decrease WNT signaling in a dose-dependent manner. **(F)** Across a range of IWP-2 concentrations, organite pattern is largely unperturbed, despite significant decreases in nuclear β-catenin. **(G)** Radial intensity profiles of normalized mean integrated Sox9 intensities. **(H)** Radial intensity profiles of normalized mean integrated Twist intensities. **(I)** DAPI, Sox9, and Twist in 24 hour, 30 hour, and 45 hour culture MO micromasses (from left to right). **(J)** DAPI, Ncad, and FN in micromass cultures (left set of images) and results from micromass simulations, ω = 2.3, at time points T=6.4, 7.2, and 8 (dimensionless units), showing distributions of cells, cell-cell adhesions, and ECM (right set of images). **(K)** DAPI, Sox9, and Twist in 24 hour WNT-treated MO micromass culture. **(L)** DAPI, Ncad, and FN in 24 hour WNT-treated micromasses (left) and results from micromass simulations with ω = 4 at time point T = 6.4 (right) exhibit accelerated pattern formation. Scale bars are 100 µm. All error bars represent mean ± SD.

Under these conditions, our continuum model spatiotemporally reproduces the dynamics of structural rearrangement into the distinct adjacent supracellular regimes observed in the organite. Specifically, our model indicates that biasing cells proximal to a central epithelial hub towards cell-cell relations causes the local ECM to recede outward to the emerging cartilage zone, where cell-ECM engagement is high and self-amplifying (Figures 6B-6C). Simultaneously, a shift from ECM engagement to cell-cell adhesion in cells adjacent to the epithelium leads to an increase in cell packing, which increases the propensity of adjacent cells to adopt cell-cell adhesion. We also confirm that inverting the organite geometry in the model reproduces the *in vivo* structural arrangement of dense ECM at the core and densely packed cells relating through cell-cell adhesion at the periphery (Figure 6D), as we observed in the limb (Figure 1A).

Our model predicts that secreted ectodermal cues need only influence cells immediately adjacent to the ectoderm (Figures 6B-6C). This local bias, coupled with the mesenchymal cross-scale co-constitution, is sufficient to generate a soft tissue domain without any long-range diffusion of an epithelial signal. Further, our model indicates that the length scale of the pattern, measured as the distance from the ectoderm to the boundary between compartments, is insensitive to a wide range of Wnt levels (Figure 6B-6C). To experimentally test the insensitivity of tissue scale patterning to Wnt levels, we pharmacologically generated a range of Wnt secretion levels^111^, confirming significant reduction of Wnt signaling in the mesenchyme via nuclear β-catenin (Figures 6E-6F, S6A). Across the range of inhibition, the pattern length scale was not significantly changed, in line with our model’s predictions (Figures 6F-6H, S6B-S6C). We observed that Wnt inhibition resulted in a modest reduction of proliferation, suggesting that epithelial Wnt increases proliferation (Figures S6D-S6G). However, given that we do not observe an effect on pattern formation at this inhibitor concentration, we conclude that this effect of Wnt on proliferation is not essential for the pattern-forming process. Taken together, our model and experiments indicate that, when the cross-scale co-constitutive capacity of mesenchyme is taken into account, patterning processes can be robust to a wide range of Wnt levels. Wnt thus biases the spatial orientation of the pattern, but the intrinsic self-organizing capacity of the mesenchymal field appears to govern the soft tissue-cartilage bifurcation itself.

We investigated this self-bifurcating potential in micromass culture, a classic, highly dense, mesenchyme-only system where such bifurcation has been observed^41-45,112-113^ and confirmed mutually exclusive Sox9 and Twist domains after 45 hours (Figure 6I). While previously observed^45^, this bifurcation has not been conceptualized or formally modeled as a process based on structural coupling across scales, leaving the structural dynamics during pattern initiation largely unexplored. We observed supracellular structure that coincided with the emergence of a Sox9 and Twist pattern. Soft tissue structure, marked by higher cell density, cell-cell adhesion, and Twist, emerged despite the absence of ectoderm (Figure 6J). We adapted our continuum model to test whether it could predict these dynamics in micromass culture. Indeed, our simulations recapitulated the observed patterns of cell density and ECM structure (Figure 6J). In both micromass and our model simulations, while an ECM network forms outside of the aggregates, we also observed central FN foci within these aggregates, consistent with residual matrix from the aggregation process and in an organization that resembles the organite (Supplemental Note and Figure S6H). The agreement between our model and experiment further supports the conclusion that the mesenchyme possesses an intrinsic, cross-scale co-constitutive capacity to bifurcate into soft tissue and cartilage fated tissue domains, independent of any patterning input from the ectoderm.

Since the micromass system establishes that mesenchyme can bifurcate without any epithelial input, it also offers a way to isolate how Wnt acts on this self-organizing capacity when reintroduced uniformly, without the spatial bias normally provided by a localized ectodermal source. To do so, we treated micromass cultures with Wnt, finding that treatment across the dense culture accelerates tissue differentiation, with domain delineation occurring within 24 rather than 45 hours (Figures 6K-6L). Incorporating this effect into simulations, by increasing the propensity of all cells for cell-cell adhesion at the expense of cell-ECM engagement, produced a similar acceleration and a more pronounced differentiation into compartments with distinct supracellular structures (Figure 6L). These results provide an example of how a signal like Wnt can alter the pattern-forming process when its levels change, not by shifting a diffusion-based gradient that encodes molecular positional information, but by tuning the structural coupling governing the mesenchyme’s intrinsic self-organizing dynamics (Figure 5L).

Taken together, our experimental results, coupled with simulations of our mathematical model, indicate that the spatial emergence of two distinct compartments arises through self-amplifying but mutually opposed co-constitutions rooted in cell-cell or cell-ECM relations. Antagonism between the two competing modes allows the mesenchyme to self-compartmentalize while remaining open to input from signals such as Wnt, which can bias the instability to reliably enable soft tissue near the ectoderm.

### (7) Cell-supracellular structural relations set organ-level pattern scale

Given that mesenchymal field patterning is insensitive to the spatial concentration distribution of Wnt, we sought to determine what sets the length scale of the pattern. By pairing experiments and simulations of our mathematical model, we tested the hypothesis that the co-constitutive relationship between cell-ECM engagement and supracellular structure has a strong causal role in establishing the pattern length scale. When we modulated this co-constitutive relationship by activating or inhibiting TRPV4 channels, the distance from the ectoderm to the cartilage-fated zone significantly increased in both cases (Figures 7A-7B, S7A), revealing a non-monotonic relationship between TRPV4 activity and pattern length scale, in which deviating from an optimal level of cell-ECM engagement in either direction disrupts the pattern. Decreasing cell-ECM engagement also caused loss of the dense, aligned fibronectin network at the edge of the cartilage-fated zone, coinciding with a blurred boundary between Sox9-negative and Sox9-positive cells (Figure S7B), indicating that sharp compartmental demarcation depends on this supracellular fibronectin structure.

**Figure 7.**
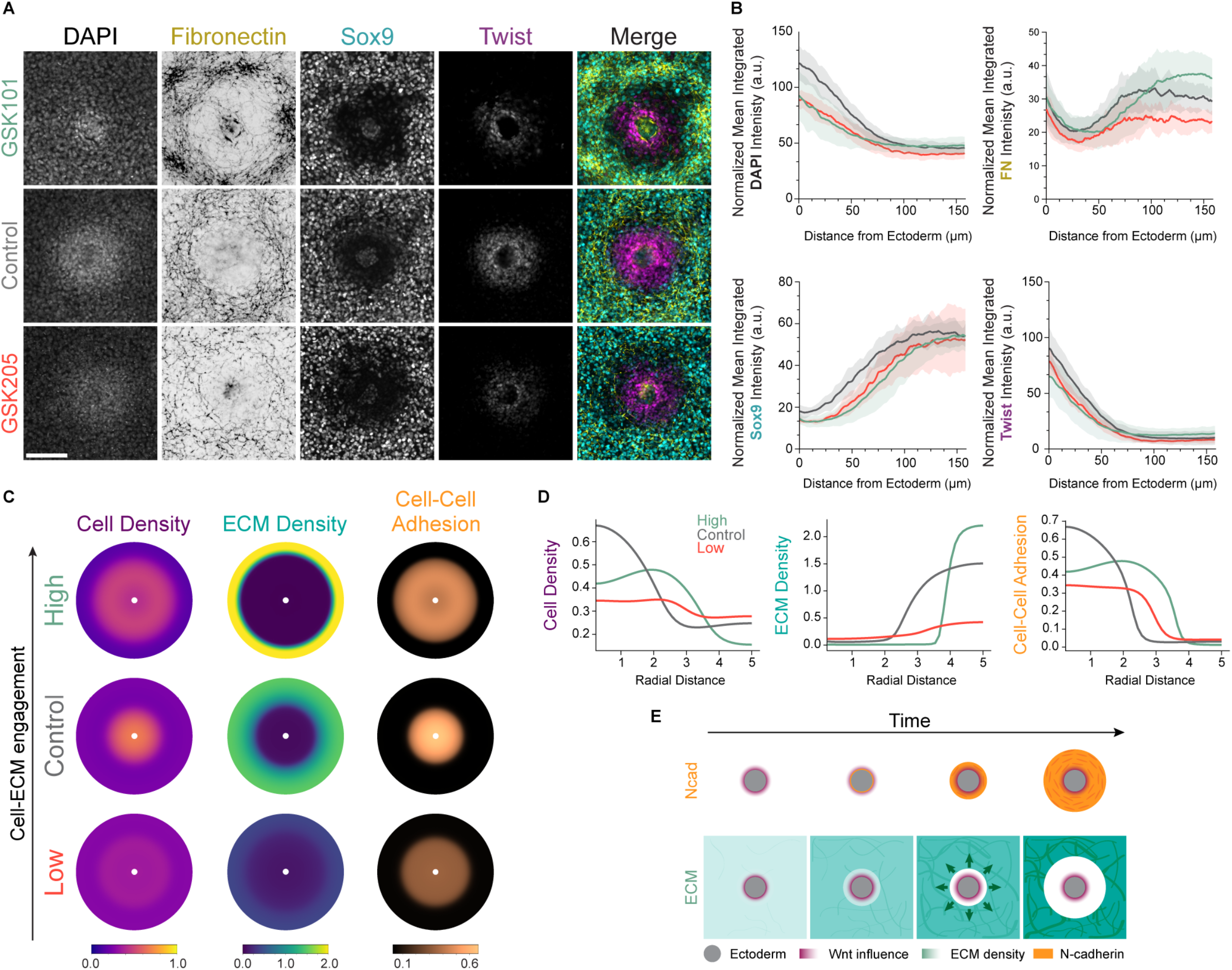
Organ-level pattern length scale depends on cell-supracellular ECM relations. **(A)** Organites treated with a TRPV4 antagonist, GSK205 (50 μM), and agonist, GSK101 (15 μM), show disrupted ECM network formation, changes in pattern length scale, and reduced Twist activation. Scale bar, 100 µm. **(B)** Mean radial intensity profiles of normalized integrated intensities of DAPI, FN, Sox9, and Twist as measured from the ectoderm. Mean ± SD shown. **(C)** Simulation snapshots at T = 4 subject to changing cell-ECM interactions showing increased ECM clearance and reduced cell density near the ectoderm when cell-ECM engagement is increased (γ: 0.5 → 1.4, Κ_b_: 2.5 → 3.1) or decreased (γ: 0.5 → 0.4, Κ_b_: 2.5 → 1.6). **(D)** Radial profiles of ECM density, cell density, and cell-cell adhesion from simulations in 7C. **(E)** Schematic summarizing the temporal dynamics of the organite pattern-forming process. Ectodermal Wnt biases cells proximal to a central epithelial hub toward cell-cell adhesion in lieu of cell-ECM engagement, causing the local ECM to recede towards the emerging cartilage zone, where cell-ECM engagement is high and self-amplifying. Simultaneously, this shift from ECM engagement to cell-cell adhesion in cells adjacent to the epithelium increases cell packing, which further increases the propensity of adjacent cells to adopt cell-cell adhesion, propagating the soft tissue zone outward until it reaches the forming cartilage’s well-developed cell-ECM network, where the cell-ECM mode is too strongly favored for cell-cell adhesion to take hold.

We observed the same effects on the pattern length scale in our simulation. Specifically, reducing or increasing the cell traction and ECM synthesis that would normally be triggered by a forming cell-ECM supracellular structure led to an increase in the distance from the ectoderm to the ECM boundary of the cartilage-fated zone in both cases (Figures 7C-7D). These results support the dependency of pattern length scale on cell-supracellular coupling through cell-ECM relations and highlight the predictive strength of our model (Figure 7E). Further, this shift in patterning and the non-monotonic effect we observe offers some rationale for why both gain- and loss-of-function TRPV4 mutations in humans are linked to skeletal dysplasias involving shortened limb skeletal segments^114-117^.

## Discussion

In this study, using a new *ex vivo* primary progenitor assay, we address how a field of limb mesenchyme transforms into distinct, adjacent cartilage- and soft tissue-fated compartments. We find that supracellular structure activates cell fate differentiation in this system. This supracellular structure is intrinsically generated through scale-crossing relations in which cell-level activity and supracellular structure reciprocally constitute their spatiotemporal co-evolution. The fact that distinct regimes of soft tissue and cartilage co-constitutions are mutually suppressive but self-amplifying transforms an initially uniform mesenchymal field into two distinct domains and also sets the resulting tissue compartment size. Lastly, epithelial Wnt enables soft tissue near the ectoderm not by providing concentration-dependent positional information to individual cells, but by locally biasing which of the two co-constitutive structural regimes takes hold.

### Cell fate specification through an intrinsically generated supracellular cue

While cell fate changes due to top-down tissue-scale effects from a neighboring tissue^25,58,92,118-119^ or extrinsic culture substrate^91,93^ have been extensively reported, here we articulate how cell fate differentiation can be stimulated through supracellular cues that progenitor cells both create and ultimately respond to by initiating either cartilage or soft tissue fate. Thus, cell fate is not the result of an externally imposed boundary condition. Prior to our study, despite extensive molecular analysis, a clear and complete explanation of how limb progenitors bifurcate into either cartilage or soft tissue fates remained elusive. An explanation based on an internally generated supracellular structural cue provides a rationale for fate specification in the context of the limb progenitor field, and may do so in other contexts where a presupposed triggering molecular ligand or external boundary condition remains elusive despite significant molecular analysis.

Furthermore, while emergent mechanical and material properties of tissues have been shown to be critical for explaining organ-level morphological changes^26,59,120-123^, our results indicate that the specific structural features that arise at the supracellular level are crucial for influencing cell structure to trigger fate changes. These results predict that to fully understand environmental influences on cell state, studies tracking features at the micro-level cell and subcellular level (e.g. individual adhesion complexes and matrix fiber) or macro-level coarse-grained mechanical properties that emerge at the tissue-scale (stiffness, viscosity) must be complemented with meso-level analyses of supracellular structural features that exist between these two scales.

### Cell-supracellular self-organization bridges molecular inputs to tissue-level phenotypes

Self-organizing systems have the capacity to internally generate greater complexity but also remain responsive and adaptive to modulation from external inputs^124^. Mechanistically characterizing tissue self-organization as a co-constitutive process based on structural coupling between the cell and supracellular level enabled us to consider how molecular inputs can be understood not as direct determinants of cell state, but rather as modulators of cross-scale structural coupling. Thus, the functional consequence of a molecular signal on internal cell state can only be understood by recognizing that cellular phenotype (propensity to adopt either cell-cell or cell-ECM relational modes) is irreducibly coupled to supracellular structural dynamics. In other organogenetic contexts, key signaling molecules may influence tissue fields not through cell-autonomous pathways that subsequently drive collective assembly, but rather by altering baseline cell-supracellular coupling dynamics.

We find that a specific organization (via cell-cell-based linkages) at the supracellular scale is necessary for cells to have the appropriate competency to respond to a Wnt signal and activate soft tissue fate. By extension, our work suggests that the functional effects of drugs, carcinogens, or environmental toxins on subcellular processes (e.g. metabolic dysfunction) may be mediated by tissue structural competency. In the context of immunological responses in which rapid changes in tissue organization can occur^125^, a full understanding of how cytokines impact stromal tissue may require uncovering cell-supracellular structural relations.

Our finding that TRPV4 modulates tissue-level patterning via cell-supracellular coupling provides a potential mechanistic inroad into understanding how genetic mutations in TRPV4 channels lead to changes in skeletal pattern^114-117^, an understanding that may be necessary for effective clinical intervention. Not accounting for cell-supracellular structural coupling may preclude a rational understanding of how certain genetic mutations cause patterning pathologies that manifest at the organ scale.

### A model for tissue symmetry breaking and patterning independent of molecular gradients

Through the identification and subsequent mathematical modeling of a self-organizing process based on cell-supracellular structural co-constitution, we provide evidence that both the initiation and development of patterns of cell fate can, in principle, occur entirely independently of morphogen concentration gradients. Traditional French Flag models assume that a key element of tissue symmetry breaking involves a spatial gradient of diffusible molecules that instruct individual cells to adopt distinct behaviors based on concentration thresholds. Rather than providing information to individual cells in a precise spatial or concentration-dependent manner, we find evidence that Wnt acts by biasing inter-level structural dynamics to seed soft tissue adjacent to the ectoderm.

The historic underlying assumption that chemical cues directly instruct individual cells to change cell fate has generally led cell and supracellular structural considerations to be subordinated in studies of molecular diffusion-based mechanisms (e.g. reaction-diffusion), especially with regard to the root causes of pattern initiation^45,126-128^. More recently, there has been increasing interest in models that merge or extend graded molecular control and mechanical processes^22-23,27,109^. The model we present here is distinct from such frameworks in that Wnt functions not by establishing a diffusion-based gradient, but by tuning the structural coupling governing the mesenchymal tissue field’s intrinsic self-organizing dynamics that occur at cell-supracellular scales.

### Cell-supracellular structural coupling as a basis for organic self-organization

Relationships between distinct scales have become increasingly appreciated in studies linking biochemical patterning with mechanical feedback from supracellular and tissue scales^17-25^. While our study also privileges cross-scale interaction, here we focus specifically on identifying and placing explanatory weight on cell and supracellular structures rather than chemical concentrations or mechanical quantities^129^. Uncovering the organizing role played by structural coupling between cell and supracellular levels allowed us to articulate previously uncharacterized functions for secreted biochemical signals in tissue patterning. It is noteworthy that at subcellular or macromolecular scales, structure is central to mechanistic explanations of causality. Our study extends this explanatory logic by showing that structures beyond the scale of a single cell can directly govern increases in tissue complexity (i.e. cell differentiation and tissue field compartmentalization).

A coupling where the constituent units are themselves transformed by their participation in the whole they build has been proposed as a defining feature of self-organization in biological hierarchies as compared to inorganic systems^34-36,130^. Furthermore, because these processes correspond to the same entity viewed at different scales, they exhibit a structural concurrence that distinguishes them from the sequential loops typical in mechanochemical feedback. There is no separation because a structural event at the cell scale (e.g., cell-ECM engagement) can inherently be linked to alteration of the supracellular structure, and a change in that supracellular structure is also linked to the structure of the cell membrane, which enables TRPV4 channel activity. Uncovering the presence and functional role of such cross-scale structural coupling provides concrete verification for the notion of concurrence, which has a strong conceptual precedent in theoretical work^30,36^.

Our finding that cell and supracellular structures require each other for initiation as well as maturation is in line with the original notion of self-organization coined by Immanuel Kant, where there is mutual determination of the part and the whole^31-32^. Recently, this notion of self-organization has been superseded by one based on a bottom-up decentralized interaction of a system of parts leading to higher order emergent properties^30,32,35^. The latter, more recent understanding allows one to draw similarities between spontaneously forming physical systems and biological systems^124,131-134^. However, the need to recover an organic notion of self-organization has been noted by biological theorists, often called “organicists” (e.g. Bertalanffy, Needham, Weiss), for over a century. Given that the relationship across levels that we observe extends beyond a merely bottom-up process to one that is co-constitutive between the part (cell) and the whole (supracellular), our study offers a novel empirically grounded case study for an organic form of self-organization specific to living systems.

## Acknowledgments

We thank members of the Laboratory of Morphogenesis at The Rockefeller University for discussion and feedback on the manuscript. We thank Drs. Ruslan Medzhitov, Cori Bargmann, and Philip Ball for comments on the manuscript and/or helpful discussions.

## Funding

Alfred P. Sloan Foundation, Innovate Award (AES)

National Institute of General Medical Sciences 1R01GM152611-02 (AES)

NIH New Innovator Award 1DP2DE033856-01 (AES)

## Author contributions

A.R.R. and A.E.S. conceived of the project together with C.S.K. A.R.R., A.E.S., and C.S.K directed the conceptual and technical development of the project. A.R.R., A.E.S., C.S.K., and R.C. designed experiments and interpreted results. C.S.K. and R.C. co-developed the organite platform, developed assays, performed experiments, and analyzed results. N.T.M. and K.C. developed assays, performed experiments, and analyzed data, with technical and conceptual guidance from C.S.K., A.R.R., and A.E.S. P.W.M. developed the mathematical model and simulations. A.R.R., A.E.S., and C.S.K. wrote the original draft of the manuscript with input and editing from R.C., N.T.M., and P.W.M. A.E.S and A.R.R co-supervised the project. All authors discussed the results and implications and commented on the manuscript at all stages.

## Data and materials availability

All data are available in the main text or the supplementary materials. Code is available here: https://zenodo.org/records/21363478

## Supplementary Materials

## Statement on AI-assisted technologies

During the preparation of this work, the author P.W.M. used GitHub Copilot for assistance in writing and debugging numerical analysis code for the mathematical model, and the author R.C. used Codex for drafting FIJI macros used for image analysis. The authors reviewed and edited the output as needed and take full responsibility for the content of the published article.

## Supplementary Materials

### Materials and Methods

#### Embryos and dissections

Fertilized chicken eggs (white leghorn) were obtained from commercial sources, incubated at 37.8°C, and staged according to Hamburger and Hamilton. GFP-labeled embryos used in this study were obtained from Clemson University and maintained in the same manner as wild-type embryos. Primary cells were obtained from embryonic day 4 forelimbs (HH21), collecting only up to the distal third of the bud and removing the posterior region to remove influences from the zone of polarizing activity. Buds were dissected in PBS and transferred to a plate where they sat in a mixture of trypsin and collagenase at room temperature for 10 min. Limb mesenchyme was manually separated from ectoderms in full media (10% fetal bovine serum, 1% penicillin-streptomycin and 2% chick serum in DMEM).

#### Cell culture assays

##### Ex vivo cultures and micromasses

Mesenchymal tissue from dissected buds (prepared as described above) were collected and mechanically dissociated with a micropipette. The suspension was passed through a 40 µm 2cell strainer (Corning) and spun down in a centrifuge at 200 rcf for 5 minutes. Supernatant was removed, and cells were resuspended in full media in the appropriate volume to produce low- or high-density suspensions. Cell suspensions were plated as 10 µL drops on standard 24-well plates for some *ex vivo* assays (Figs. S1B, 2I, 2J, 4J, S4A), Ibidi 8-well slides for organites (Cat. 80826), and15-well µ slides for micromasses and most other MO cultures (Cat. 81506) and incubated at 39°C for 2 hours before full media (with or without treatments) were added. For cultures that went longer than 24 hrs (micromasses at 30 and 45 hrs in Figure 6), media was replaced daily.

##### Organite platform

Organites were prepared in a similar fashion to our *ex vivo* or micromass cultures, except for these assays ectodermal tissue was included with the mesenchymal tissue before mechanical dissociation. Thus, rather than peeling away all ectoderms after collagenase/trypsin treatment during embryo dissections, a percentage of excised limb forelimbs were left unpeeled (20-30%). Cell suspensions of mesenchymal and epithelial tissue were plated in 10 µL drops in Ibidi 8-well chamber slides. For Fig. S1D, limb buds from GFP-labeled embryos were mixed in, typically at <5% of all limb buds.

##### Treatments

The following pharmacological agents were utilized across cell culture assays to perturb our experimental systems: FAKi (FAK inhibitor 14, Tocris, Cat. 3414), Y-27632 (Tocris, Cat. 1254), TRPV4 inhibitor (GSK205, MedChemExpress, Cat. HY-120691A), TRPV4 activator (GSK101, Selleck, Cat. S8107), L-type Ca2+ channel activator (Bay K8644, Tocris, Cat. 1544), Calmodulin inhibitor (W-7, Tocris, Cat. 0369), and WNT secretion inhibitor (IWP-2, Tocris, Cat. 3533). In addition, we utilized the following proteins and peptides as treatments across all our assays: FUD peptide (synthesized by PSL GmbH, HPLC-purified and verified), Collagenase Type 1 (Worthington, Cat. LS004196), WNT (Human Wnt-3a, Tocris, Cat. 5036-WN), Fibronectin (Recombinant human fibronectin protein, Tocris, Cat. 4305-FNB), Ncad inhibitor (ADH-1, MedChemExpress, HY-13541), and Integrin activating antibodies (TASC, Sigma-Aldrich, Cat. MAB19294; TS2/16, Invitrogen, Cat. MA2910).

#### Immunofluorescence and Imaging

##### In vitro culture assays

All samples were fixed in 4% paraformaldehyde in PBS for 20 minutes at room temperature. After washes in PBS and PBTx (0.1% Triton-X in PBS), samples were treated with CAS-Block (Invitrogen) for 30 minutes at room temperature. Once the blocking agent was removed, samples were incubated with primary antibodies diluted in 10% CAS in PBTx for 2-3 hours at room temperature or overnight at 4°C. The following primary antibodies and dilutions were used: Sox9 (Abcam, ab5535; 1:1000), Twist (Santa Cruz, sc-81417; 1:300), N-cadherin (Proteintech, 22018-1-AP; 1:300 & Sigma Aldrich, C3865; 1:300), AF488-conjugated Fibronectin (Thermo Fisher, 53-9869-80; 1:300), Paxillin (5H11) (Thermo Fisher, AHO0492; 1:300), β-catenin (Santa Cruz, sc-59737; 1:100), β1 Integrin (Sigma Aldrich, I8638, 1:300), Vinculin (Sigma Aldrich, V9131, 1:300), Thrombospondin-1 (Invitrogen, MA5-13398; 1:300), type IIa Collagen (Santa Cruz, sc-52658; 1:300), Tenascin (Thermo Fisher, MA5-16086; 1:300), and E-cadherin (Abcam, ab76055; 1:300).

After staining with primary antibodies, samples were washed with PBTx and then incubated with secondary antibodies diluted in 10% CAS in PBTx for 2-3 hours at room temperature or overnight at 4°C. The following secondary antibodies and dilutions were used: DAPI (Invitrogen, D1306, 1:1000), Alexa Fluor 488 (Invitrogen, 1:300), Alexa Fluor 555 (Invitrogen, 1:300), Alexa Fluor 647 (Invitrogen, 1:300), and ReadyProbes™ Phalloidin conjugates (Thermo Fisher, Cat. R37178, 1 drop/mL). After staining, samples were imaged on a Nikon ECLIPSE Ti2 with either a Plan Apo 4X (NA 0.2) or S Plan Fluor 20X (NA 0.45) objective. Some images (Figs. 5A, 6I and K, S6I) were imaged on a Zeiss LSM980 with a 25X objective or a Zeiss LSM780 (Figs. S1C, S2B).

##### In vivo sections

For visualization of limb tissue (Figs. 1A-C, S1A and 2H), whole embryos were fixed in 4% paraformaldehyde overnight at 4°C and then transferred to 30% sucrose in PBS for another overnight incubation. Limb buds were cut within the sucrose solution with forceps and transferred to Tissue-Tek O.C.T. compound (Sakura, Cat. 4583) and stored at -80°C. These frozen samples were sectioned with a Cryostat (Leica CM3050S) and tissue sections were collected on Superfrost slides (Fisherbrand, Cat. 12-550-15). Slides were treated with the same immunostaining protocol described above.

##### Live imaging

Both standard *ex vivo* culture assays and organites were plated in Ibidi 8-well chamber slides, and imaged in a Zeiss Celldiscoverer 7. The culture chamber was set at 37°C and images were acquired once every 6 minutes with a 20X objective lens set to 0.5x zoom. For live visualization of fibronectin in *ex vivo* assays or organites, AF488-conjugated FN antibody was added to the media (1:300 dilution). For holotomography to generate 3D stacks (Fig. S1C and Movie S1), organites plated in Ibidi 8-well chamber slides were imaged in a Tomocube HT-X1 Plus).

#### Image Analysis and quantification

Prior to analysis, all images were processed by first producing a maximum intensity projection with all z slices that were in focus (typically 2 slices). Some images were additionally processed with background subtraction (see below).

Analysis of DAPI, Sox9, Twist, and FN profiles in the organite platform was performed using the Radial Profile Angle FIJI plugin (https://imagej.net/ij/plugins/radial-profile.html). The radius was set at 600 pixels (Figs. 7 and S7), 500 pixels (Fig. 1D), 475 pixels (Figs. S6E-F), 450 pixels (Figs. 5I-J), 400 pixels (Fig. 5N), or 380 pixels (Figure 6), where the circular region was placed with the center of circle aligned with the center of the ectodermal tissue. Radial profiles were calculated on the entire stack for all replicates. To remove confounding signals measured from within the ectodermal tissue, a cutoff was established based on the Twist profile and removed from the dataset. The data were then recalibrated such that the x axis values represent distance from the edge of the ectoderm rather than the center of the circular ROI. Radial intensity profiles of DAPI, Sox9, and Twist were normalized for Fig. 1D to show relative profiles for all channels on the same plot. For Fig. 5J, the radial intensity profile for the expected increase in Twist intensity at higher cell densities was produced by first measuring the average % increase in DAPI intensity as a function of distance from the ectoderm. These values were then multiplied to the Twist intensity profiles of each replicate. Each radial line intensity profile was integrated through an area under the curve (AUC) analysis in GraphPad Prism, which was calculated for all replicates across all conditions. These values were compared for statistical analyses (see Statistical Analysis).

Cell aspect ratios in the organite platform were measured in experiments where a low percentage (<5%) of equivalently-staged limb mesenchyme from GFP-labeled chicks were mixed in. Images were processed through thresholding and basic watershed transformation before individual GFP cells were identified through Analyze Particles where measurements like cell aspect ratio were made.

For the EdU analysis (Figs. S6D – G), the Click-iT™ EdU Cell Proliferation Kit (Invitrogen, Cat. C10340) was performed at roughly 22 hours into organite culture. Samples were fixed at 24 hours and stained for DAPI following standard immunostaining protocols. Images were analyzed to generate radial intensity profiles of DAPI and EdU Signal (as described above). EdU profiles for each sample were divided by their respective DAPI profiles to normalize for cell density. Linear regression analysis was performed in GraphPad Prism.

Analysis of DAPI, Sox9, Twist, and FN profiles in *ex vivo* assays were performed using Fiji (ImageJ, NIH). For transcription factor expression (Sox9 and Twist), the Sox9 or Twist channel images were converted into a binary mask via thresholding to measure the total Sox9^+^/Twist^+^ area. DAPI images were converted into a binary mask and measured for total nuclear area. The percentage of Sox9/Twist^+^ area was calculated as: % Sox9^+^/Twist^+^ area = (Sox9^+^/Twist^+^ mask area / DAPI mask area) × 100. Since DAPI staining was consistent with high signal-to-noise ratios across samples, background subtraction was not applied to the DAPI channel. All values were normalized to the percentage of Sox9/Twist^+^ area of the control group into Sox9/Twist expression fold change. Fibronectin intensity (Figs. 2O, 3D, 4D, 4I, S4E) was quantified by measuring the integrated density of the Fibronectin channel. Background subtraction was not applied as the fibronectin staining was consistent with high signal-to-noise ratios across samples. N-cadherin^+^ area (Figs. S3B, 5B, 5E) was measured from binary masks created from thresholding. For Fig. 5C, N-cadherin^+^ area normalized to DAPI was calculated as (N-cadherin^+^ area/DAPI^+^ area) × 100.

The histograms of fibronectin fiber orientations were generated with the Directionality FIJI plugin, utilizing Fourier component analysis on 200 x 400 pixel regions that were oriented parallel to the circumferential axis of the fibronectin ring. For Fig. S7B, the same ROIs used to produce the orientation histograms were also measured for mean FN intensities in ImageJ. The comparison to the cartilage network outside of the ring was made with the same ROIs for rings but placed further within the cartilage tissue.

#### Statistical Analysis

See Supplementary Table S1 for full documentation of experimental trials and sample sizes. Statistical comparisons between two groups were performed using unpaired Mann– Whitney tests in GraphPad Prism. All tests were two-tailed and non-parametric. Error bars represent the standard deviation (SD), and exact p-values and sample sizes (n) are reported in the figure legends. For radial intensity profile comparison across conditions, the area under the curve (AUC) of each replicate’s profile was calculated and used as a single-value summary. A Kruskal–Wallis test was used to assess statistical significance across conditions.

#### Simulations

A mathematical model was designed to help evaluate some of our hypotheses regarding supracellular patterning. We constructed a nonlinear system of partial differential equations that represent the essential mechanical factors that we suspect are organizing both cells and matrix in our experiments. The full details of this model are presented in the Supplementary Text; however, we provide a brief summary of its structure here. The experimental system is represented as a series of continuum fields interacting in a bounded 2D domain (either a disk, an annulus or a periodic box). The patterning of cells is described by their density *c*(*x, t*), which evolves via a combination of diffusion and cell-cell adhesion. The ECM is treated as a compressible viscoelastic fluid, characterized by its density *ρ*(*x, t*), bulk pressure *p*(*x, t*), and deviatoric stress ***σ***(x, t). Material is constantly added to the matrix in proportion to the local cell density, and cells likewise deform the matrix through active contractility. Finally, we characterize the coupling between cellular behavior and the supracellular environment by introducing two feedback variables *I*(*x, t*) and *N*(*x, t*), which represent the mean commitment of cells at (*x, t*) to cell-ECM or cell-cell interactions, respectively. We introduce mutual repression between these two variables, leading cells to favor one or the other class of interaction in a way that is modulated by local cell and ECM density. These variables are co-moving with the local cell population and so are also subject to advection and diffusion.

## Supplemental Material

A structured-population continuum model for cell-supracellular-based self-organization of a mesenchymal tissue field

### 1 Introduction

This document presents a continuum model for mesenchymal tissue self-organization that couples four interacting components: cell-extrinsic density evolution, cell-extrinsic extracellular matrix (ECM) mechanics, cell-intrinsic phenotype switching, and ectoderm-derived signaling. The model is intentionally minimal: each term is included because it is needed to capture a specific experimentally observed behavior, such as aggregation, matrix accumulation, fate-dependent feedback, or boundary-driven spatial bias. In this sense, the framework is designed as a mechanistic bridge between cell-level interpretations and tissue-scale observables (domain-level pattern wavelength, ring structure, and radial localization).

Relative to classical limb-patterning models based on Turing-type activator-inhibitor reaction-diffusion systems [10, 11], our organizing principle is different. Spatial structure is driven primarily by cell-extrinsic collective migration and cell-extrinsic ECM remodeling, represented here by Cahn-Hilliard-type cell-density dynamics coupled to active viscoelastic matrix flow. This framing is connected to the modelling lineage of Murray, Oster, and Harris [6], but extends that class of models by explicitly including a co-evolving internal cell state: the variables *N* and *I* describe local commitment to cell-cell versus cell-ECM engagement. That addition places the model in the broader category of structured-population or internal-state continuum descriptions [12].

For clarity, the distinction from classical Turing models can be summarized in three points. First, pattern initiation does not come from a finite-wavelength activator-inhibitor linear instability; instead it emerges from aggregation/coarsening-like cell dynamics (spinodal/binodal phase-separation-like behavior) coupled to matrix mechanics. Second, the model is intrinsically nonequilibrium because active stress and continuous ECM production (source term *k*_*b*_*Ic*) inject energy and mass over time, so the system is not relaxing toward a closed thermodynamic steady state. Third, transport is not diffusion-only: advection, mechanically induced fluxes, and viscoelastic matrix flow are central, so the governing dynamics are fundamentally beyond classical reaction-diffusion form.

The material below is organized as follows. Section 2 outlines the key aspects of our model, providing commentary on specific assumptions and modeling choices. Section 3 describes the rescaling and non-dimensionalization of our problem. Section 4 then outlines our numerical scheme.

### 2 Governing Equations

#### 2.1 Variables of interest

Our model consists of the following fields over a bounded two-dimensional spatial domain Ω. In the following sections, we define the governing equations for these fields and explain the associated modeling assumptions.

| Cell-extrinsic |  |
| --- | --- |
| $c(\mathbf{x}, t)$ | Cell density |
| $\mu(\mathbf{x}, t)$ | Chemical potential |
| $\rho(\mathbf{x}, t)$ | ECM density |
| $\mathbf{v}(\mathbf{x}, t)$ | ECM velocity |
| $p(\mathbf{x}, t)$ | ECM pressure field |
| $\boldsymbol{\sigma}(\mathbf{x}, t)$ | ECM deviatoric stress tensor |
| Cell-intrinsic |  |
| $I(\mathbf{x}, t)$ | Integrin commitment |
| $N(\mathbf{x}, t)$ | N-cadherin commitment |

#### 2.2 Cell Density and Chemical Potential

In our model, cells are treated as a conserved scalar field that evolves through linear diffusion and several competing interactions. We represent these interactions through an effective chemical potential, *µ*, for convenience and consistency with the FEM scheme described below. We include two complementary aggregation mechanisms: a cell-cell adhesion term based on local cell density and mediated by N-cadherin (*N*), and an ECM-mediated interaction term controlled by *I*. We also include a quadratic repulsive potential to represent short-range steric effects in crowded environments, along with a gradient penalty term that suppresses short-wavelength instabilities at interfaces. We neglect cell division based on experimental evidence suggesting that proliferation plays a limited role on the timescales considered here. Taken together, the cell-density equations are:

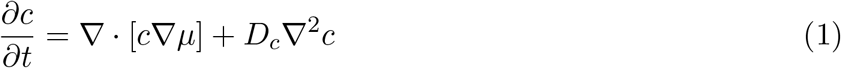

where chemical potential is given by:

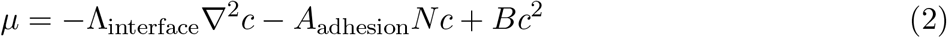

and

- *D*_*c*_ is a cell diffusion rate
- *A*_adhesion_ is the cell-cell adhesion coefficient
- Λ_interface_ is an interfacial energy cost
- *B* is a coefficient restricting cell crowding

In the absence of ECM (*ρ* = 0), the long-timescale dynamics reduce to standard binodal phase separation between high- and low-density cell regions.

#### 2.3 ECM evolution

For this model, we adopt a deliberately simplified description of the extracellular matrix as an isotropic, passive viscoelastic fluid. This choice improves interpretability and computational tractability, but it necessarily sacrifices fidelity to the true ECM constitutive behavior, which reflects a complex cross-linked network. More accurate coarse-grained theories of active cross-linked networks remain an open theoretical problem beyond the scope of this work [4]. Despite this simplification, the present constitutive law captures the key trends observed experimentally.

A key feature of our problem, in contrast to previous models of cell-ECM interaction [8], is the presence of large variations in matrix density. Consequently, we first introduce a scalar ECM density *ρ* which obeys a continuity equation:

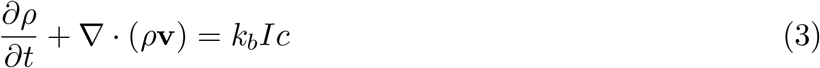

Consistent with experiments showing monotonic ECM accumulation over time, we include a cell-dependent matrix synthesis term with rate *k*_*b*_ and intentionally omit degradation. As a result, the full system generally lacks a steady state, except in the special case *Ic* = 0 everywhere. ECM velocity *v* is determined from force balance, with contributions from baseline viscosity, bulk elastic pressure *p*, shear stress ***σ***, and an isotropic active contractile stress:

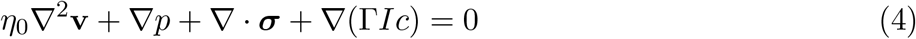

Here,

- *η*_0_ is a small basal ECM viscosity
- Γ*>* 0 is a cell-ECM traction parameter establishing a contractile active force

We assume that the elastic component of the matrix follows Maxwell-type dynamics with bulk modulus *K*, shear modulus *G*, and relaxation timescale *T*_relax_. The resulting bulk-pressure evolution is

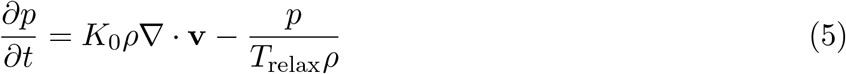

and the stress tensor as:

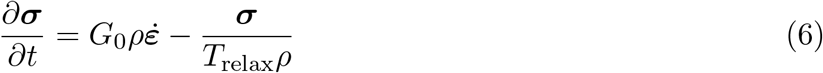

Here 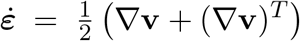. We assume density-dependent constitutive parameters to capture density-induced stiffening. In this model, increasing ECM density both raises the elastic moduli and lengthens stress-relaxation times, so denser ECM behaves as both stiffer and more solid-like.

#### 2.4 Cell Behavior Dynamics

An essential feature of our model is mutual inhibition between cell-cell and cell-ECM engagement modes. We use a simple resource-allocation picture: each cell has a finite pool of cytoskeletal machinery, so increased commitment to one mode reduces commitment to the other. We represent these commitments with *N* (cell-cell, N-cadherin-associated) and *I* (cell-ECM, integrin-associated), and model their competition as a local bistable switch. The inhibition strength is modulated by the local supracellular environment: high ECM density weakens repression of the cell-ECM mode, whereas high cell density and high Wnt exposure weaken repression of the cell-cell mode.

|  |  |  |
| --- | --- | --- |
| Cell density | $c_0$ | $\sqrt{\frac{D_c}{b}}$ |
| Length | $L_0$ | $\left(\frac{\Lambda_{\text{interface}}}{D_c b}\right)^{1/4}$ |
| Time | $T_0$ | $\left(\frac{\Lambda_{\text{interface}}^2}{D_c^3 b}\right)^{1/2}$ |
| ECM density | $\rho_0$ | $\left(\frac{\Lambda_{\text{interface}}^2}{D_c^3 b T_{\text{relax}}^2}\right)^{1/2}$ |
| Stress | $\sigma_0$ | $\frac{K}{T_{\text{relax}}^2} \left(\frac{\Lambda_{\text{interface}}^2}{D_c^3 b}\right)^{1/2}$ |
| Wnt | $L_W$ | $\frac{1}{k_w} \left(\frac{\Lambda_{\text{interface}}^2}{D_c^3 b T_{\text{relax}}^2}\right)^{1/2}$ |

Transport is also essential. Cell-intrinsic dynamics should be interpreted in a co-moving frame with the cells rather than purely in the Eulerian lab frame (see Ref. [9]). Accordingly, we include transport terms that account for both bulk cell motion and diffusive mixing. The resulting equations of motion are:

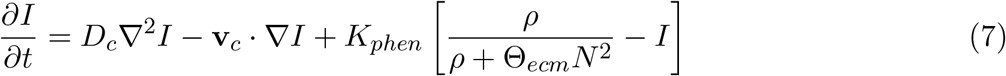

and

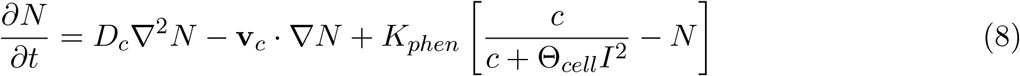

where:

- *K*_*phen*_ is the phenotype switching rate
- Θ_*ecm*_ is the ECM threshold parameter
- Θ_*cell*_ is the cell threshold parameter
- **v**_*c*_ = − ∇ *µ -D*_*c*_∇ log *c* is the density-normalized cell flux

### 3 Rescaling

To simplify the model, we rescale variables using the following characteristic scales:

#### 3.1 Complete System

After nondimensionalization, the complete governing system is:

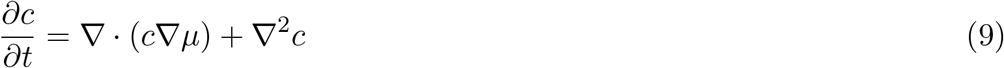

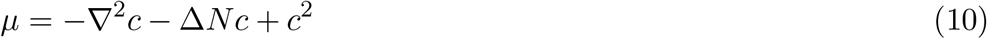

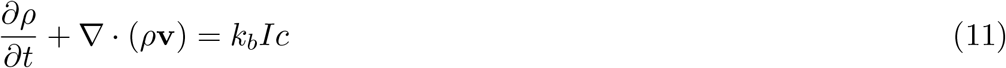

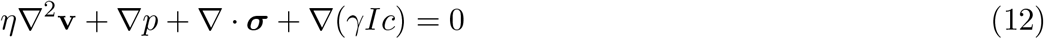

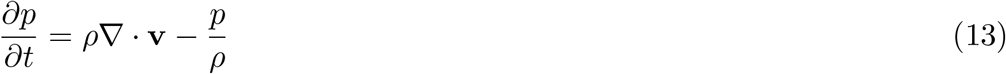

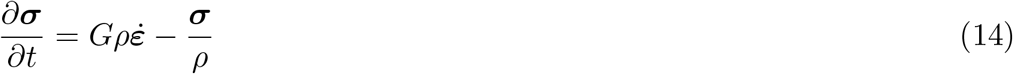

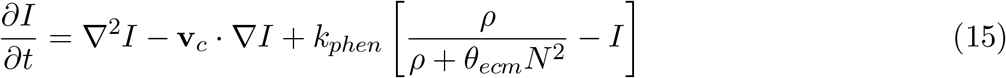

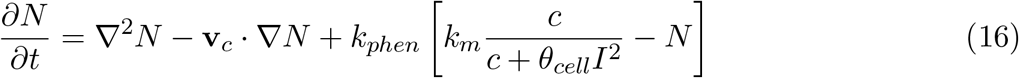

#### 3.2 Boundary Conditions

On annular and disk domains, the boundary conditions for the system are:

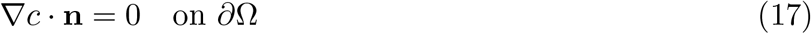

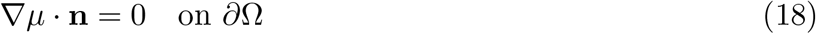

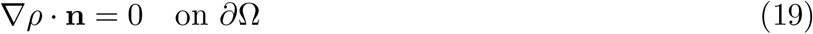

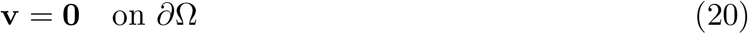

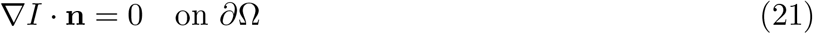

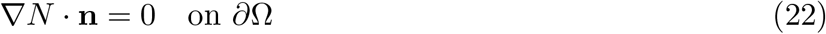

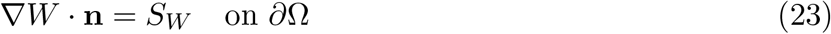

Here, *S*_*W*_ is zero except on the inner boundary of the annular simulations where it serves as a source term for *W* .

#### 3.3 Initial Conditions

In all scenarios, we used initial conditions of the form:

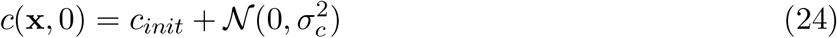

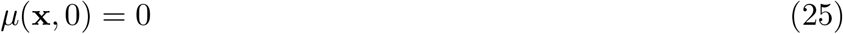

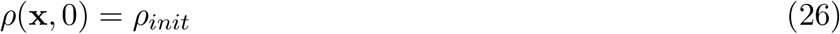

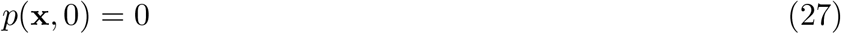

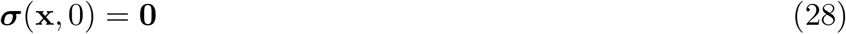

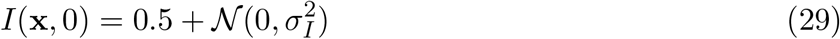

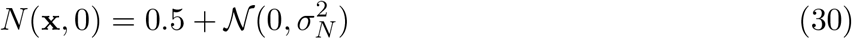

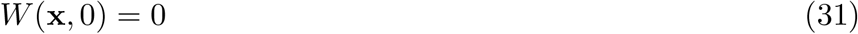

For organite and inverted simulations, we used *σ*_*c*_ = 10^*-*4^ and *σ*_*N*_ = *σ*_*I*_ = 10^*-*6^. For the purely mesenchyme scenario, we instead set *σ*_*c*_ = 0.18 and *σ*_*N*_ = *σ*_*I*_ = 10^*-*8^. The substantially larger cell-density noise reflects a qualitative estimate that initial cell distributions in mesenchyme-only assays are relatively heterogeneous. This perturbation amplitude was chosen to provide a reasonable match to the observed timescales in mesenchyme-only and organite variants of the system. To prevent negative initial values, we threshold perturbed cell densities so that *c* ≥ 0.01 everywhere.

### 4 Perturbations

#### Mesenchyme-only micromass

In this scenario, we model the effect of Wnt dosage by introducing a global bias that upregulates *N*, leading to increased cell aggregation. Specifically, we modify the equation of motion to

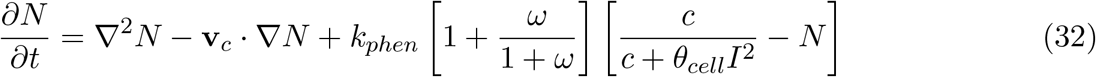

where the scalar constant *!* is taken to be a proxy for Wnt dosage.

#### Organite and inverted scenarios

##### Ectodermal signal

By contrast, to represent the spatially localized effect of the ectoderm, we introduce a scalar field *W* (*x*) obtained by solving

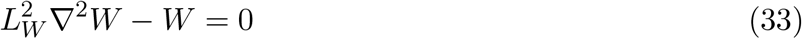

subject to a constant-flux boundary condition at the ectodermal boundary and no-flux conditions elsewhere. This field can be interpreted as a proxy for Wnt signaling, together with other ectodermal effects such as MMP secretion that favor cell-cell engagement. It modulates the cell-intrinsic dynamics as

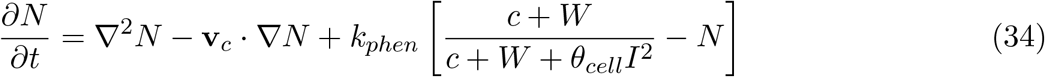

and

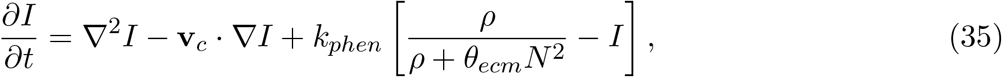

effectively increasing the threshold of ECM required to repress cell-cell interaction. Ectodermal perturbations were implemented by tuning the strength of this source term.

##### ECM Engagement

The active contractility, and synthesis rate *k*_*b*_ govern ECM organization in this model. We represented perturbations of cell-ECM engagement by jointly increasing or decreasing these two parameters, corresponding to reduced or enhanced ability of cells to form focal-adhesion-mediated interactions with the ECM.

**Table 1:**
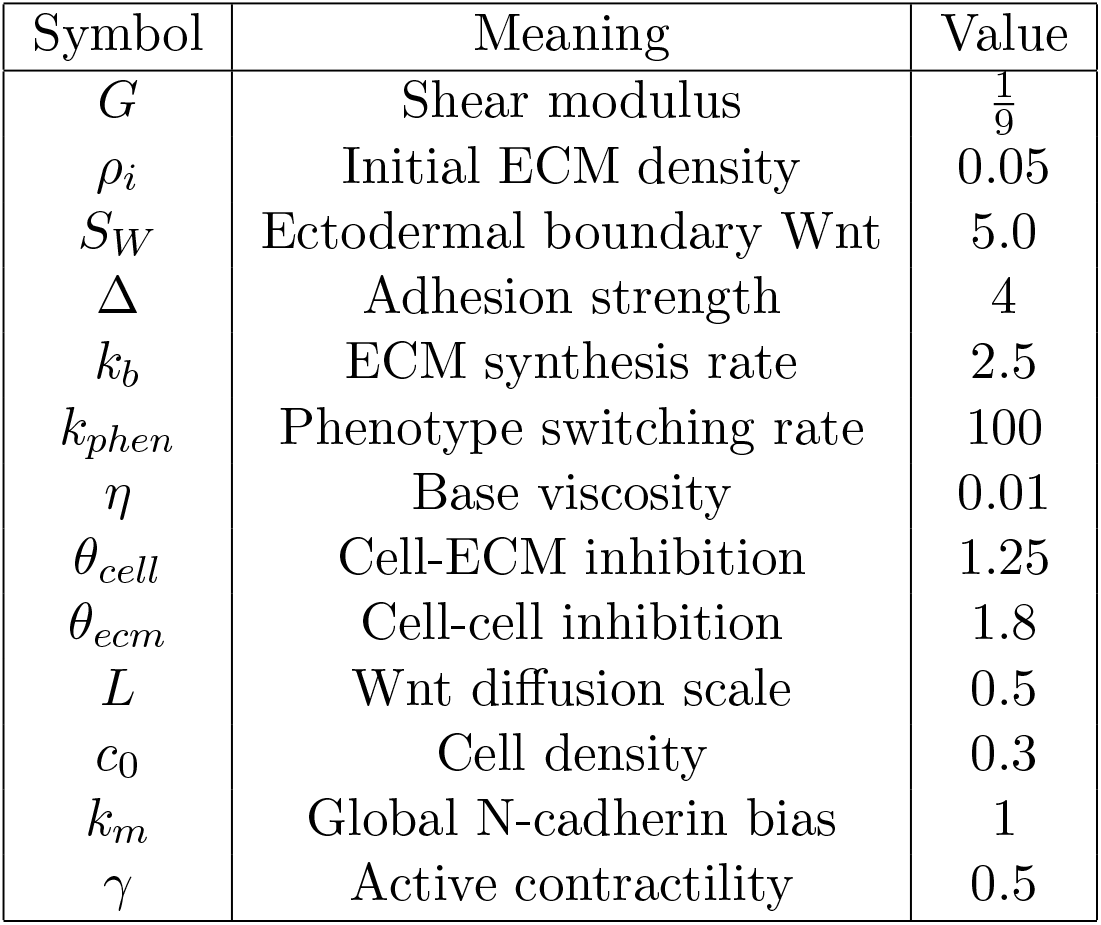
Baseline parameter values.

| Symbol | Meaning | Value |
| --- | --- | --- |
| $G$ | Shear modulus | $\frac{1}{9}$ |
| $\rho_i$ | Initial ECM density | 0.05 |
| $S_W$ | Ectodermal boundary Wnt | 5.0 |
| $\Delta$ | Adhesion strength | 4 |
| $k_b$ | ECM synthesis rate | 2.5 |
| $k_{phen}$ | Phenotype switching rate | 100 |
| $\eta$ | Base viscosity | 0.01 |
| $\theta_{cell}$ | Cell-ECM inhibition | 1.25 |
| $\theta_{ecm}$ | Cell-cell inhibition | 1.8 |
| $L$ | Wnt diffusion scale | 0.5 |
| $c_0$ | Cell density | 0.3 |
| $k_m$ | Global N-cadherin bias | 1 |
| $\gamma$ | Active contractility | 0.5 |

### 5 Numerical Implementation Notes

We employed two numerical schemes to solve the systems above. For the organite and inverted scenarios on annular and disk domains, we used finite elements on a triangulated mesh. For mesenchyme-only micromass simulations, a spectral approach was more efficient. Both schemes are described below, and code is available on request.

#### 5.1 FEM Scheme

The weak-form time-stepping scheme uses a semi-implicit treatment in which linear terms are implicit and nonlinear terms are explicit. We split the time-stepping problem into several matrix blocks that are solved in sequence. Within each block, we use a mixed finite element formulation and solve the resulting linear systems with PETSc Krylov solvers. We outline the weak form below, using (·, ·) for the *L*^2^ inner product over Ω and (·, ·)_∂Ω_ for the boundary integral. We denote by *ϕ*_*c*_, *ϕ*_*µ*_, *ϕ*_*ρ*_, *ϕ*_*v*_, *ϕ*_*p*_,, *ϕ*_*I*_, *ϕ*_*N*_ test functions from the appropriate first-order Lagrange finite element spaces. The cell-density subproblem is then:

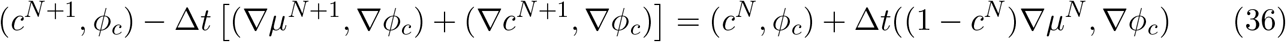

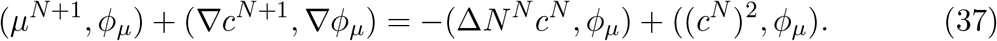

This scheme introduces a small artificial viscosity of order Δ*t* in the time evolution of *c*, which improves stability under strong advective fluxes at modest cost to accuracy. The ECM subproblem is:

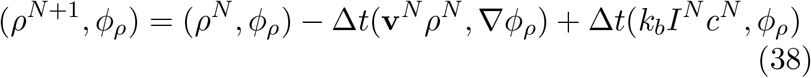

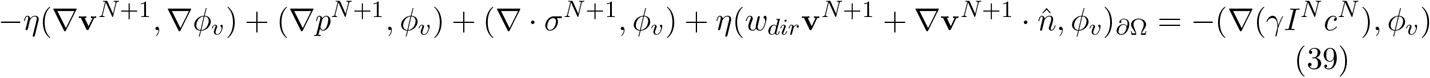

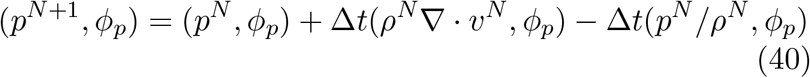

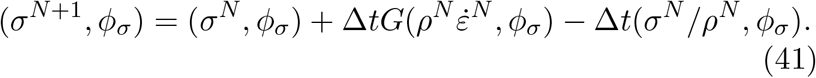

We impose the Dirichlet boundary condition on *v* weakly, with penalty weight *w*_*dir*_ = 10^3^. The cell-phenotype equations are discretized as

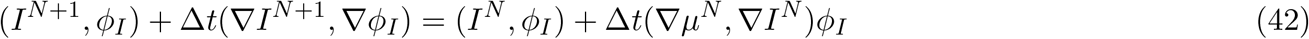

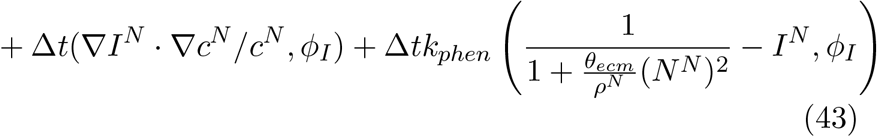

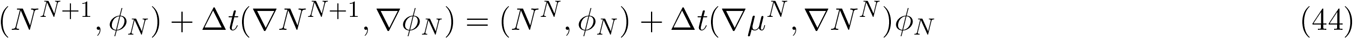

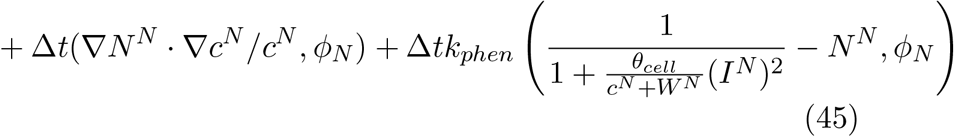

Finally, we calculate the Wnt field by solving

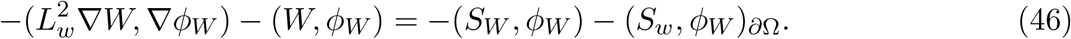

The spatial discretization uses first-order finite elements for all fields. The implementation uses the Dolfinx FEM library with PETSc linear solvers [2, 1]. Time integration is performed using a first-order semi-implicit scheme, with fixed timestep Δ*t* = 1 × 10^*-*3^. In all cases, the outer domain radius was *R*_*out*_ = 5, and for annuli, the inner boundary was *R*_*in*_ = 0.25.

#### 5.2 Spectral Scheme

In the micromass periodic-domain scenario, a spectral scheme was advantageous for two reasons. First, standard conforming finite elements are prone to instability under the large advective ECM fluxes in this regime. Second, the large periodic box (*L*_*x*_ = *L*_*y*_ = 20) is handled efficiently in Fourier space. We therefore performed these simulations with the Dedalus spectral PDE package [**?**], using real-Fourier modes and operator splitting to update the ECM, cell, and phenotype subproblems at each timestep. Time integration uses the built-in SBDF2 scheme. Relative to the FEM approach, we additionally included numerical diffusivity in the ECM mass-transport equation for stability:

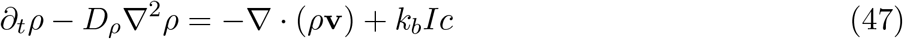

where *D*_*ρ*_ = 10^*-*4^ was sufficiently small to have a negligible effect on the simulation results.

---

**Supplemental Figure S1.**
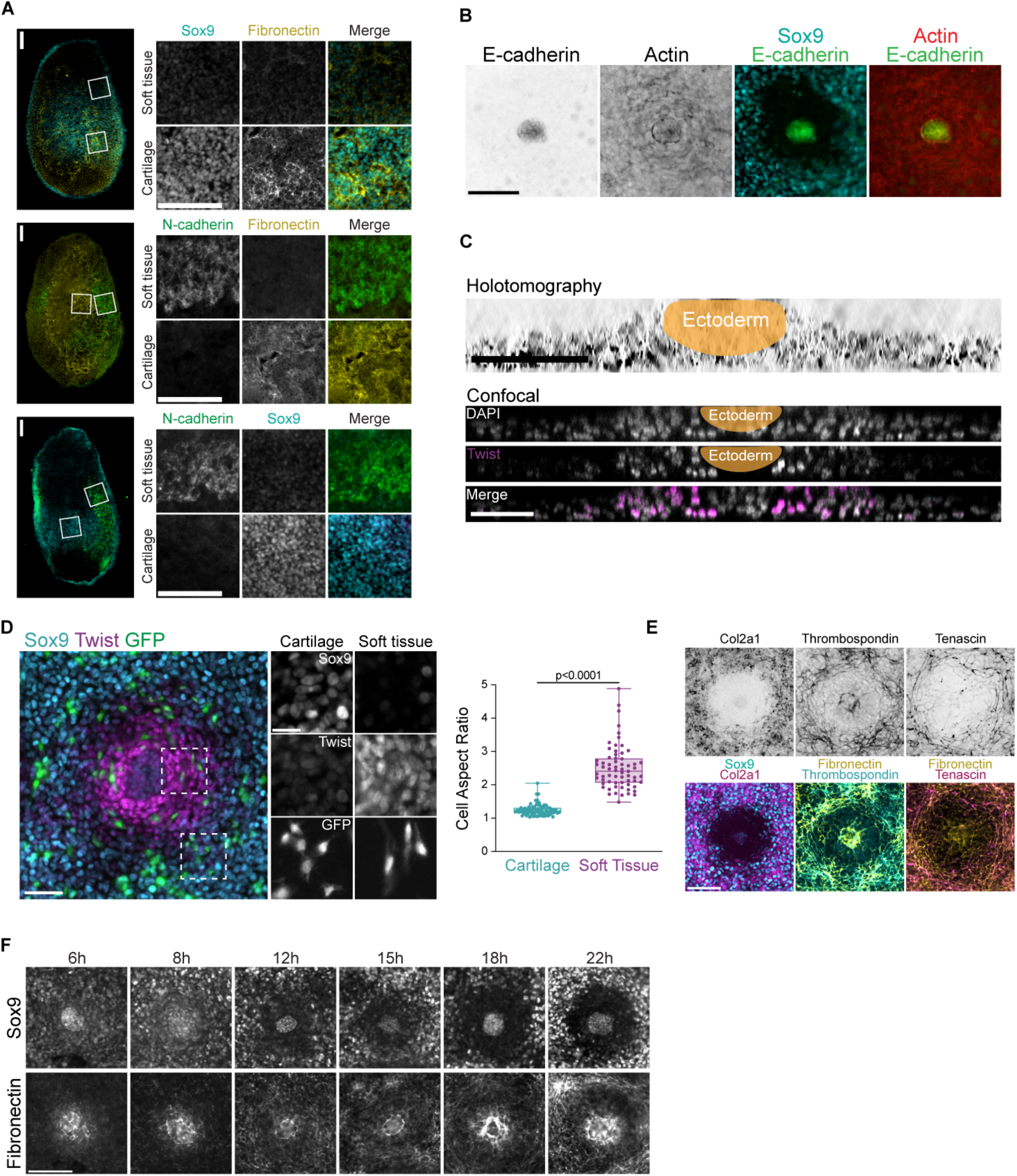
**(A)** Sections of E4.5 limb buds (left) stained for Sox9 and FN (top), Ncad and FN (middle), and Ncad and Sox9 (bottom). Crops of regions in white boxes are shown on the right. **(B)** Organite stained for E-cad, Sox9, and actin to confirm the presence of epithelial ectoderm. **(C)** Orthogonal views from z-stacks generated by holotomography (top) or confocal microscopy (bottom) demonstrating organite culture is a multi-cell thick disc. Confocal images stained for DAPI and Twist. Scale bars, 50 µm. **(D)** Left, Sox9 and Twist in an organite with <5% GFP cells reveal soft tissue cells are more elongated than cartilage cells. Scale bars, 50 μm (left) and 20 μm (right). Right, quantification of cell aspect ratios comparing cartilage and soft tissue. **(E)** Organites stained for type IIa collagen, thrombospondin, tenascin, Sox9, and FN. **(F)** Time course of organites fixed and stained for **FN** and Sox9 indicating early expression of FN in the ectoderm and across the mesenchyme. Scale bars are 100 µm unless otherwise noted.

**Supplemental Figure S2.**
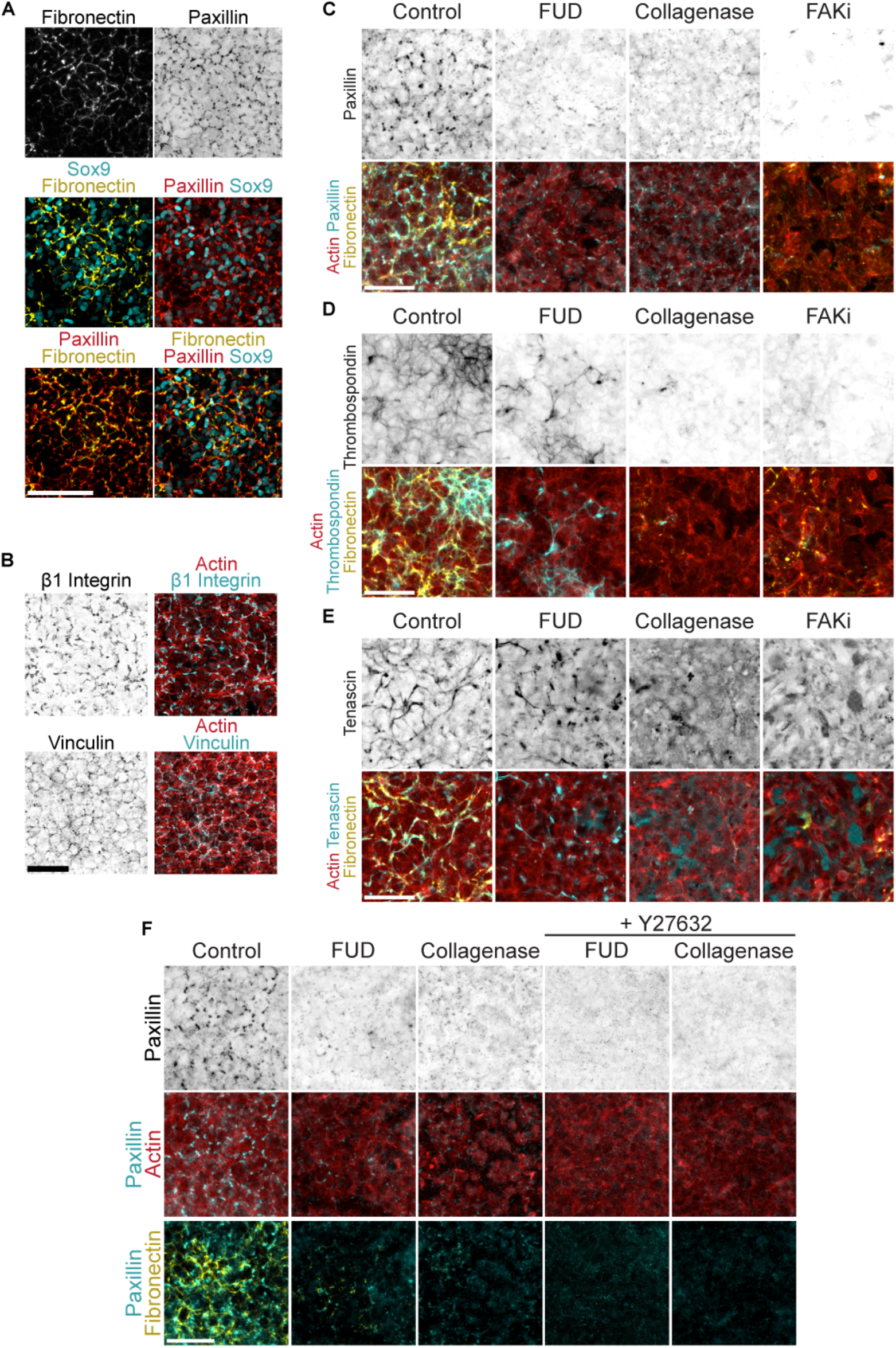
**(A)** Focal adhesion components are enriched at interfaces between cells in 24-hour *ex vivo* cultures. MO cultures stained for FN, paxillin, and Sox9. Scale bar, 100 µm. **(B)** DAPI, actin, β_1_ integrin, and vinculin in MO cultures. **(C)** MO cultures treated with FUD (0.5 µM), collagenase (0.5 µg/mL), and FAKi (20 µM) stained for paxillin, actin, and FN. **(D)** Same perturbations as S2C, stained for thrombospondin, actin, and FN. **(E)** Same perturbations as S2C-D, stained for tenascin, actin, and FN. **(F)** Paxillin, actin, and Sox9 in 24-hour MO cultures treated with FUD, collagenase, and Y27632 (10 µM). Scale bars are 50 µm unless otherwise noted.

**Supplemental Figure S3.**
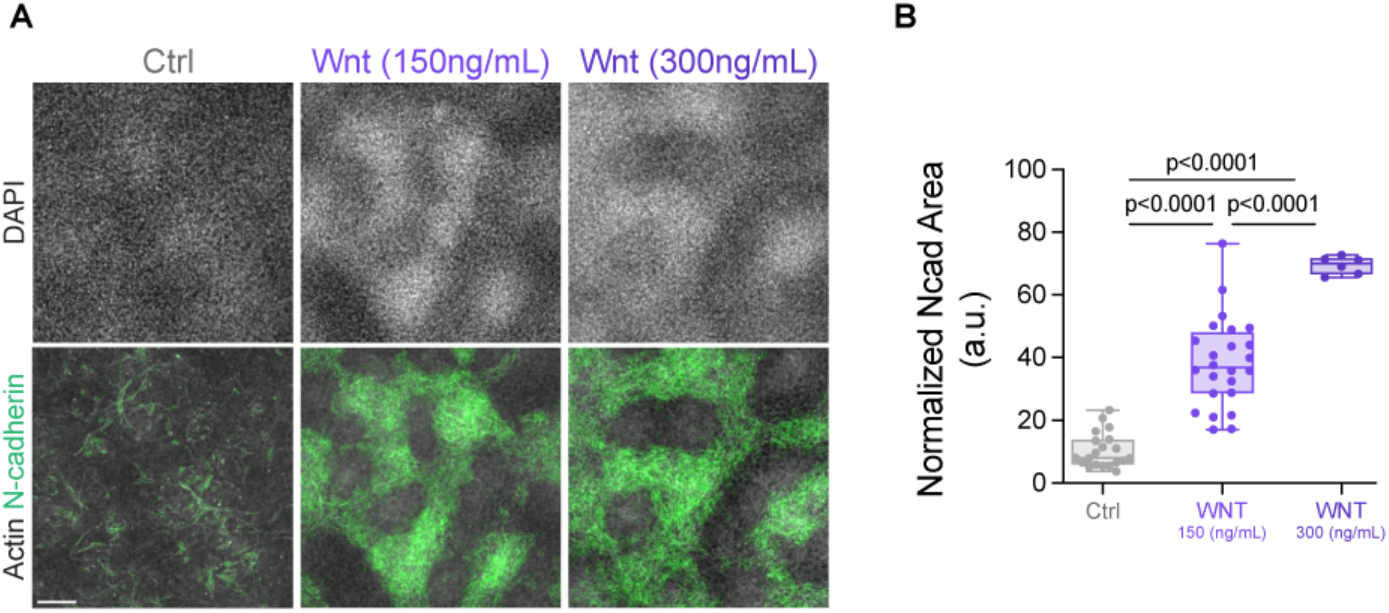
**(A)** MO cultures treated with 150 or 300 ng/mL WNT stained for DAPI, Nead, and actin show WNT induces Nead and cell packing in a dose-dependent manner. Scale bar, 100 µm. **(B)** Quantification of normalized Nead area.

**Supplemental Figure S4.**
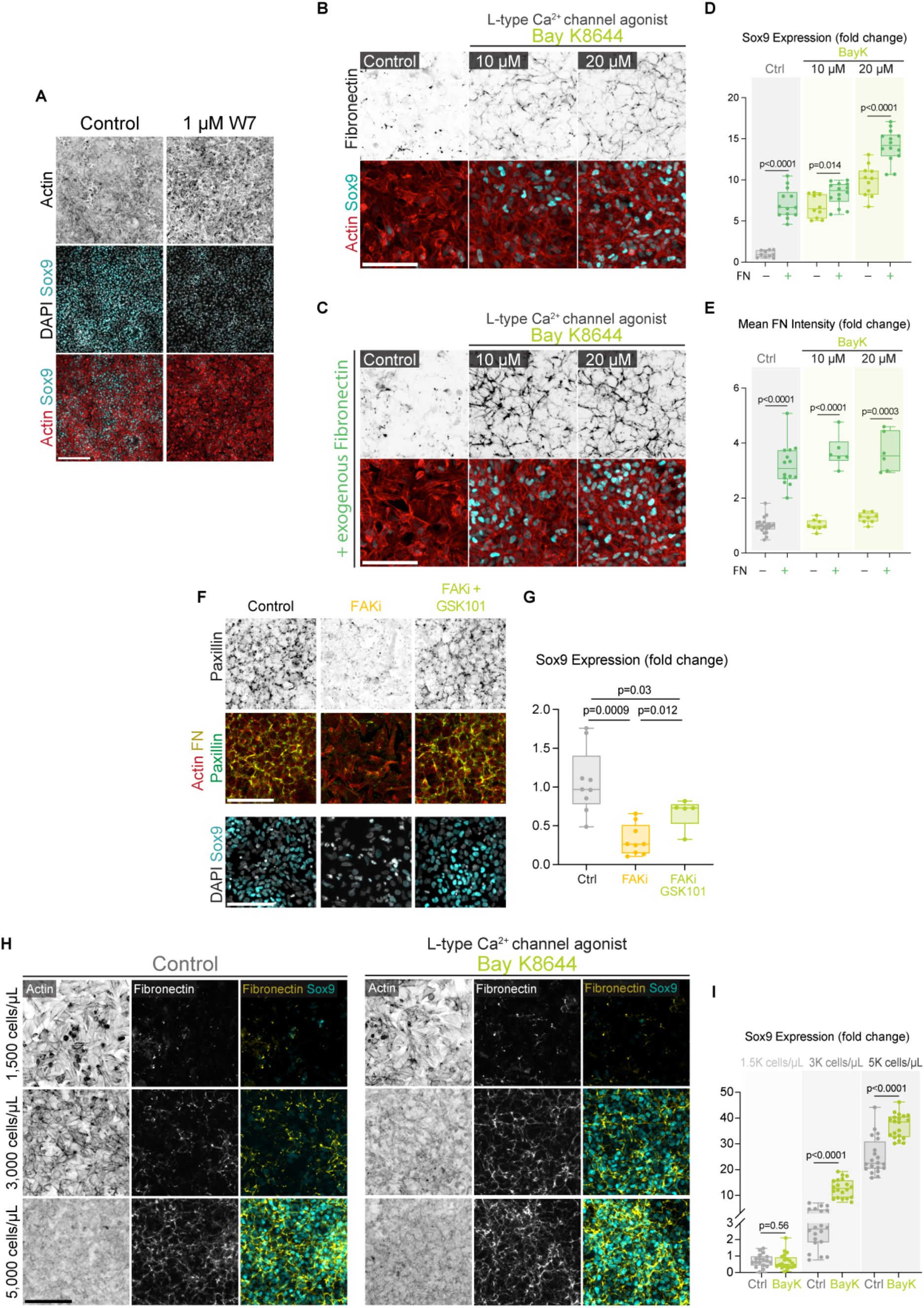
**(A)** DAPI, Sox9, and Actin in MO cultures treated with a calmodulin inhibitor, W-7 (1 µM). Scale bar, 200 µm. **(B)** Medium-low density MO cultures stained for FN, actin, and Sox9 treated with increasing concentrations of the L-type Ca^2+^ channel agonist Bay K8644 (10 – 20µM). **(C)** Same culture conditions as S4B but with the addition of exogenous FN. **(D)** Quantification of Sox9 levels (fold change) for S4B and S4C. **(E)** Quantification of FN intensity (fold change) for S4B and S4C. **(F)** Activation of TRPV4 (15 µM GSK101) rescues cell-ECM engagement and Sox9 in MO cultures treated with FAKi (20 µM). Cultures were stained for DAPI, Sox9, paxillin, actin, and FN. Sox9 images are from a separate replicate. **(G)** Quantification of changes in Sox9 levels. **(H)** MO cultures plated at a range of cell densities (1,500 to 5,000 cells/µL) and treated with Bay K8644 (20 µM) stained for actin, FN, and Sox9. **(I)** Quantification of changes in Sox9 levels in S4H. Scale bars are 100 µm unless otherwise noted.

**Supplemental Figure S5.**
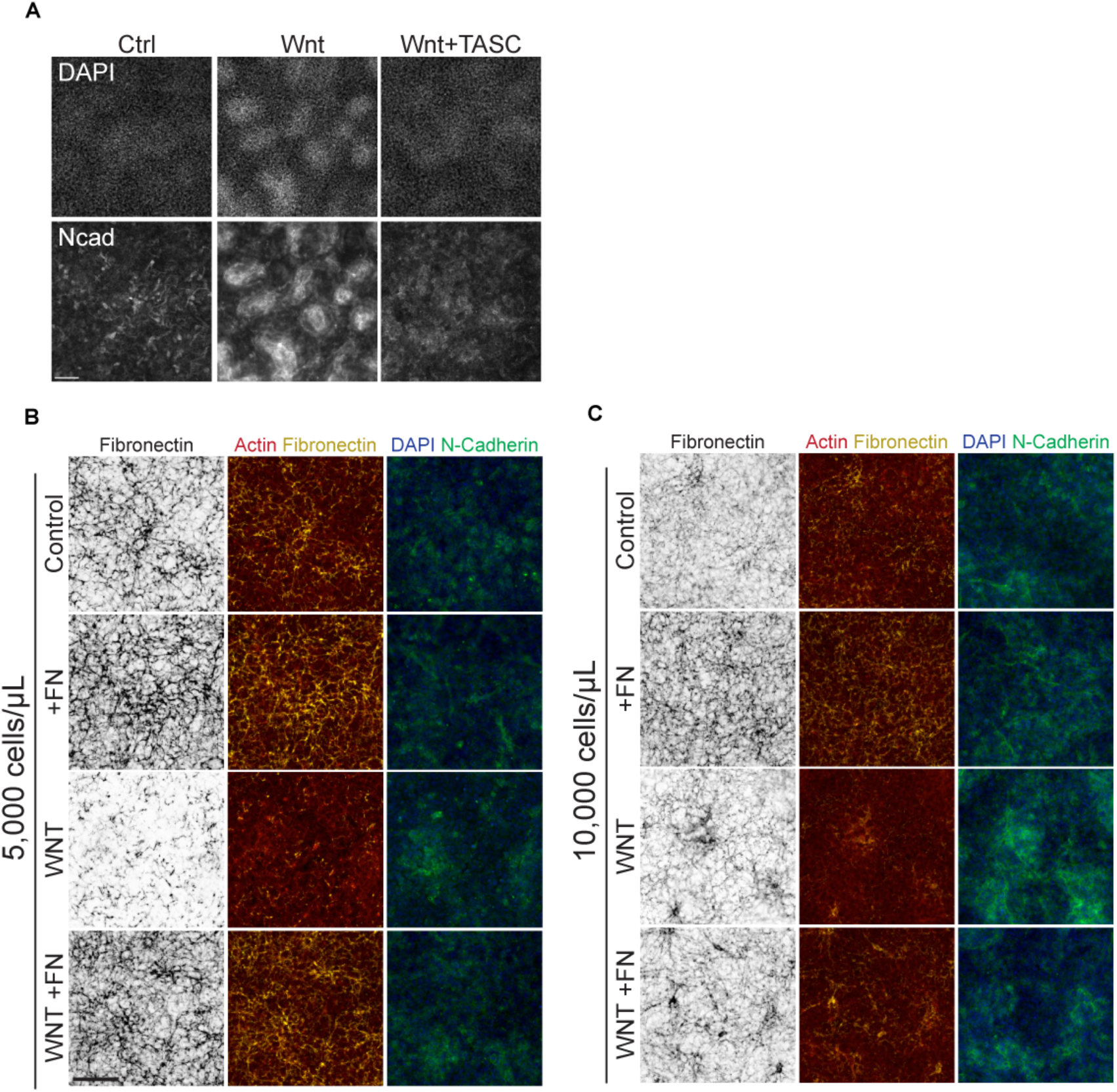
**(A)** DAPI and Nead in control or WNT-treated (150 ng/mL) MO cultures of limb progenitors with PI integrin activation (TASC, 20 µg/mL) show decreased Nead when cell-ECM engagement is increased. **(B)** Medium-high density MO cultures (5,000 cells/µL) treated with WNT (150 ng/mL), with or without exogenous FN (5 µg/mL). **(C)** Increasing density (10,000 cells/µL) and Nead in WNT-treated MO cultures decreased the degree to which exogenous FN was incorporated. Scale bars are 100 µm unless otherwise noted.

**Supplemental Figure S6.**
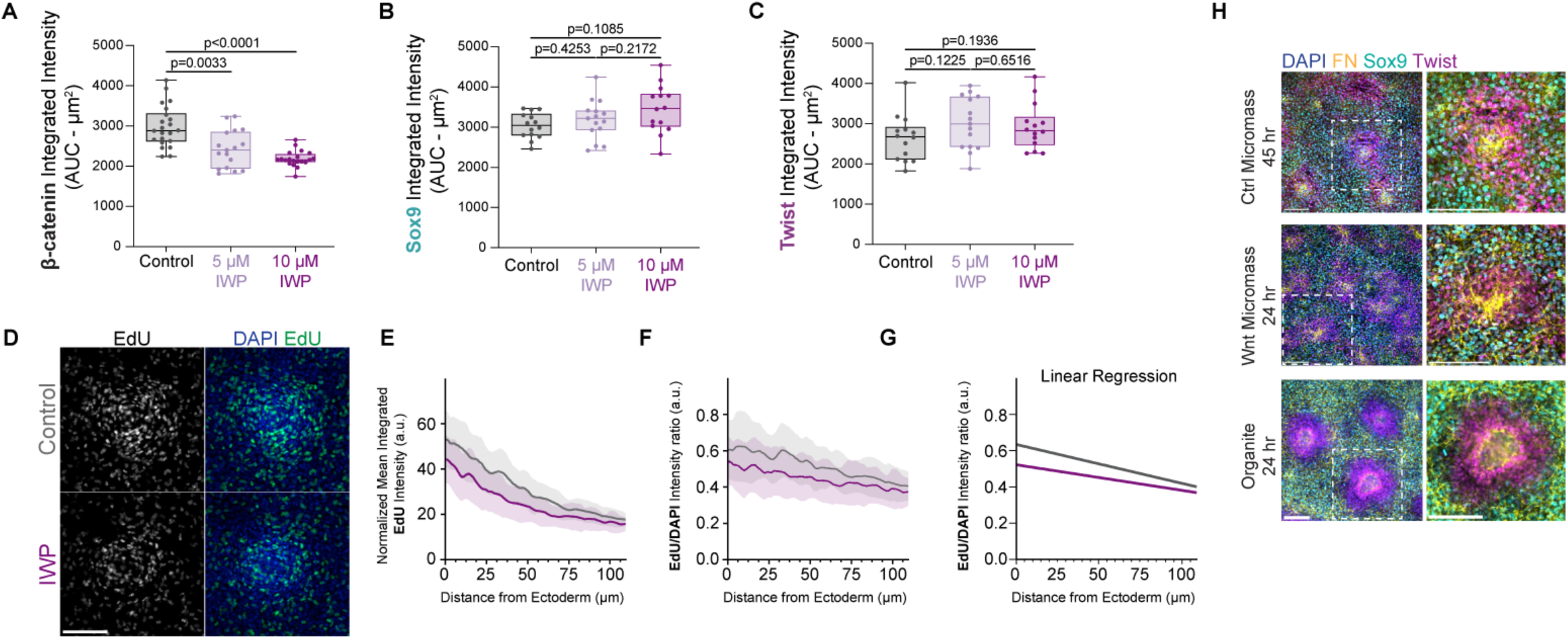
**(A)** AUC analysis of normalized mean integrated intensities of β-catenin from Figure 6D. **(B)** AUC analysis of normalized mean integrated intensities of Sox9 from Figure 6F. **(C)** AUC analysis of normalized mean integrated intensities of Twist from Figure 6G. **(D)** Control and IWP-2-treated (10µM) organites at 24 hours stained for DAPI and EdU (added at 22 hours of culture). **(E)** Radial intensity profiles of normalized mean integrated intensities of EdU. **(F)** Radial intensity profiles of normalized mean integrated intensities of EdU normalized to DAPI (EdU/DAPI ratio). **(G)** Linear regression analysis of S6G. Control y-intercept is 0.635 and IWP-2 y-intercept is 0.523 (p=0.0161). Slopes are not significantly different. **(H)** Merged images of DAPI, fibronectin, Sox9, and Twist in a 45-hour control micromass (top) and a 24-hour micromass treated with Wnt (middle) show similarities in organization to organite culture (bottom). Scale bars are 100 µm. All error bars represent mean± SD.

**Figure S7.**
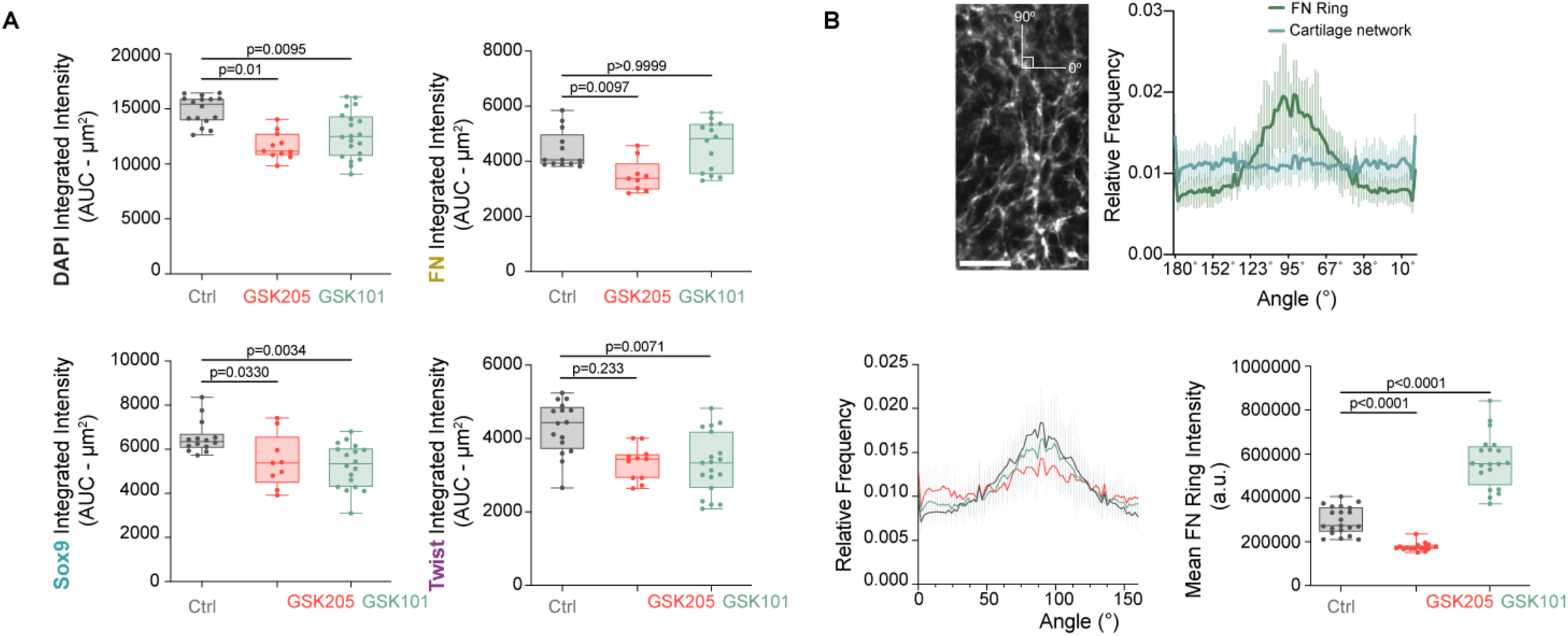
**(A)** Comparison of integrated intensities through an area under the curve (AUC) analysis of plots in 5D shows significant disruptions in patterns when TRPV4 is either inhibited or activated. All error bars represent mean± SD. **(B)** (Top) Histogram of FN fiber orientation in the FN boundary compared to the non-boundary areas of cartilage. Fiber orientations were measured with the boundary’s circumferential axis parallel to 90° (see left image). Scale bar, 25 µm. (Bottom, left) Histogram of fibronectin fiber orientation angles within the FN ring immediately neighboring the soft tissue. A χ^2^ test was performed comparing each treatment group to control: GSK205, p <.0001; GSK101, ns. (Bottom, right) Comparison of mean integrated intensities of FN.

**Table S1.** Statistical Values.

| Experiment/Figure | Trials | Total n analyzed |
| --- | --- | --- |
| Fig. 1D (organite radial intensity profile) | 5 | 12 |
| Fig. S1D (cell aspect ratios) | 2 | 78 cartilage cells, 65 soft tissue cells |
| Fig. 2C (%Sox9 time course) | 3 | 8 per time point |
| Fig. 2D (%Sox9 across density) | 3 | 23 for 1k cells/ μL, 20 for 1.5k cells/ μL, 16 for 5k cells/ μL |
| Fig. 2G (Sox9 expression with ECM perturbation) | 2 | 12 control, 8 FUD, 13 collagenase, 12 FAKi |
| Fig. 2K (Sox9 expression with Y27632) | 3 | 24 control, 20 Y27632 |
| Fig. 2N (Sox9 expression with ECM perturbations and rescue with Y27632) | 2 | 20 control, 20 FUD, 20 Collagenase, 20 FUD+Y27632, 11 Collagenase+Y27632 |
| Fig. 2O (Mean FN intensity with ECM perturbations and rescue with Y27632) | 2 | 20 control, 20 FUD, 10 Collagenase, 20 FUD+Y27632, 11 Collagenase+Y27632 |
| Fig. 3D (Exogeneous FN supplement to control and WNT micromasses) | 2 | 12 per condition (control-control, control-FN, WNT-control, WNT-FN) |
| Fig. 3F (Exogeneous FN supplement to control and WNT micromasses) | 2 | 16 control-control, 13 control-FN, 7 WNT-control, 17 WNT-FN) |
| Fig. 3J (Twist expression Wnt and ADH-1 treatment to micromasses) | 3 | 8 per condition (Control, WNT, WNT+ADH-1) |
| Fig. 3H (Sox9 expression with WNT and integrin activating perturbations) | 2 | 19 control, 20 WNT, 17 WNT+TASC, 17 Wnt+TS2/16 |
| Fig. S3B (N-cadherin expression with different WNT concentrations) | 2 | 19 control, 24 WNT (150ng/mL), 6 WNT (300ng/mL) |
| Fig. 4B (Sox9 expression with BAPTA-AM and exogenous fibronectin) | 4 | 24 control-control, 22 control-FN, 23 BAPTA(10μM)-control, 18 BAPTA(10μM)-FN, 22 BAPTA(20μM)-control, 18 BAPTA(20μM)-FN, 22 BAPTA(40μM)-control, 20 BAPTA(20μM)-FN. |
| Fig. 4D (Mean FN intensity with BAPTA-AM and exogenous fibronectin) | 4 | 24 control-control, 21 control-FN, 23 BAPTA(10μM)-control, 18 BAPTA(10μM)-FN, 22 BAPTA(20μM)-control, 18 BAPTA(20μM)-FN, 22 BAPTA(40μM)-control, 20 BAPTA(20μM)-FN. |
| Fig. 4H (Sox9 expression with GSK101 and exogenous fibronectin) | 2 | 10 control-control, 13 control-FN, 13 GSK101(10μM)-control, 14 GSK101(10μM)-FN, 13 GSK101(20μM)-control, 13 GSK101(20μM)-FN. |
| Fig. 4I (Mean FN intensity with GSK101 and exogenous fibronectin) | 2 | 12 control-control, 14 control-FN, 10 GSK101(10μM)-control, 14 GSK101(10μM)-FN, 10 GSK101(20μM)-control, 14 GSK101(20μM)-FN. |
| Fig 4K (Sox9 expression with Wnt and GSK101 treatment) | 2 | 21 control, 16 WNT, 24 WNT+GSK101 |
| Fig 4M (GSK101 treatment across densities) | 2 | 18 control-1.5k cells/μL, 20 control-3k cells/μL, 19 control-5k cells/μL, 19 GSK101-1.5k cells/μL, 20 GSK101-3k cells/μL, 20 GSK101-5k cells/μL. |
| Fig. S4D (Sox9 expression with Bay K8644 and exogenous fibronectin) | 2 | 10 control-control, 13 control-FN, 10 BayK(10μM)-control, 14 BayK (10μM)-FN, 11 BayK (20μM)-control, 14 BayK (20μM)-FN. |
| Fig. S4E (Mean FN intensity with Bay K8644 and exogenous fibronectin) | 2 | 21 control-control, 14 control-FN, 8 BayK(10μM)-control, 6 BayK (10μM)-FN, 8 BayK (20μM)-control, 6 BayK (20μM)-FN. |
| Fig. S4G (Sox9 expression with FAKi and GSK101) | 2 | 9 control, 9 FAKi, 5 FAKi+GSK101 |
| Fig S4I (Bay K8644 treatment across densities) | 2 | 18 control-1.5k cells/μL, 20 control-3k cells/μL, 20 control-5k cells/μL, 20 GSK101-1.5k cells/μL, 20 GSK101-3k cells/μL, 20 GSK101-5k cells/μL. |
| Fig. 5E (Ncad expression with WNT and ADH-1 across densities) | 2 | 8 control, 8 WNT, 8 WNT+ADH-1, 12 WNT+ADH-1 at high density |
| Fig. 5G (Twist expression across densities) | 2 | 8 per condition (1k cells/μL, 1.5k cells/μL, 5k cells/μL) |
| Fig. 5I-K (Twist intensity comparing low vs. high density organites) | 3 | 8 low density, 9 high density |
| Fig. 5N (Ncad levels after TASC treatment in organites) | 4 | 21 per condition |
| Fig. 6E, S6A (β-catenin profile after IWP-2 treatment) | 5 | 24 control, 19 5μM IWP-2, 20 10μM IWP-2 |
| Fig. 6G-H, Fig. S6B-C (Sox9 and Twist profile after IWP-2 treatment) | 4 | 14 control, 15 5μM IWP-2, 14 10μM IWP-2 |
| Fig. S6E-G (EdU analysis) | 3 | 25 control, 25 IWP-2 |
| Fig. 7B, Fig. S7A (Profile analysis in organites) | 7 | 16 control, 11 GSK205, 20 GSK101 |
| Fig. S7B (FN orientation in ring vs. cartilage) | 7 | 17 ring measurements, 21 non-ring measurements |
| Fig. S7B (FN orientation in treatment groups) | 5 | 21 control, 16 GSK205, 20 GSK101 |

## Supplemental Videos

**Movie S1. Z-stack of an organite with holotomography**. Organite cultured in an incubated Tomocube HT-X1. Imaging begins from the slide surface and moves up towards the ectoderm. Movie is running at 21 fps. Scale bar, 100 µm.

**Movie S2. Sparse labeling with GFP+ cells reveal flattening and elongation of cells in the soft tissue**. (Left) GFP signal only; (Right) Merge with brightfield. Organites were prepped with ∼5% GFP cells (see Methods) and imaged on a Zeiss CD7. Fast moving GFP-positive cells are loose cells crawling on top of the culture. Movie is running at 21 fps. Scale bar, 100 µm.

**Movie S3. Low ECM in the soft tissue of the organite system emerges through a dynamic remodeling process**. Organites were cultured with AF488-conjugated FN antibodies on a Zeiss CD7. Movie is running run at 21 fps. Time stamps are displayed as hours:minutes. Scale bar, 100 µm.

**Movie S4. Clearing of FN associated with outwards “flow” of fibers away from the ectoderm**. Organites were cultured with AF488-conjugated FN antibodies on a Zeiss CD7. Movie is running at 21 fps. Time stamps are displayed as hours:minutes. Scale bar, 100 µm.

**Movie S5. Fibronectin (FN) increases in density to form a dense network in limb progenitor *ex vivo* cultures**. *Ex vivo* MO culture treated with AF488-conjugated FN antibodies (FN-3) imaged on a Zeiss CD7. Movie is running at 21 fps. Time stamps are displayed as hours:minutes. Scale bar, 100 µm.

**Movie S6. FN network dynamics in MO cultures treated with Wnt**. *Ex vivo* MO cultures treated with Wnt and incubated with AF488-conjugated FN antibodies imaged on a Zeiss CD7. Movie is running at 21 fps. Time stamps are displayed as hours:minutes. Scale bar, 100 µm.

